# Deciphering the epitranscriptomic code of RNA degradation with nanopore direct RNA sequencing

**DOI:** 10.64898/2026.05.09.723953

**Authors:** Zhang Zhang, Chen-Long Wang, Zhi-Hao Zhang, Yi-Feng Huang, Ying-Yuan Xie, Zhen-Dong Zhong, Guo-Run Tang, Ze-Hui Ren, Ye-Lin Lan, Jin-Wen Kong, Zhi-Shuo Qiao, Tao-Wen Su, Hong-Xuan Chen, Qiu-Yu Wang, Ru-Jia Luo, Jun-Tong He, Wen-Qing Liu, Fu Wu, Guan-Zheng Luo

**Affiliations:** State Key Laboratory of Biocontrol, MOE Key Laboratory of Gene Function and Regulation, Guangdong Province Key Laboratory of Pharmaceutical Functional Genes, School of Life Sciences, Sun Yat-sen University, Guangzhou 510275, China; Sun Yat-sen University Institute of Advanced Studies Hong Kong, Science Park, Hong Kong SAR, 999077, China; Innovation Center for Evolutionary Synthetic Biology, Sun Yat-sen University, Guangzhou 510275, China

## Abstract

The precise regulation of RNA degradation is crucial for gene expression homeostasis, yet how multiple molecular features coordinate on a single transcript remains poorly understood. Here, we use nanopore direct RNA sequencing (DRS) to simultaneously track alternative isoforms, m6A modifications, and poly(A) tail dynamics at single-molecule resolution during a time course of RNA decay. We show that m6A regulates RNA degradation in a stoichiometry-dependent manner, where modification levels quantitatively modulate decay kinetics. Mechanistically, m6A is functionally coupled to deadenylation, promoting accelerated poly(A) tail shortening and coordinated RNA turnover. At the isoform level, we identify regional m6A clusters (RMCs) as structural elements that associate with isoform-selective degradation and remodel protein-coding potential. Furthermore, transcript splicing architecture is associated with distinct m6A deposition patterns, suggesting that gene structure encodes RNA decay kinetics through m6A-mediated regulation. A machine learning model integrating these multi-modal features highlights the central contribution of m6A and deadenylation in shaping RNA decay, while revealing substantial regulatory heterogeneity across transcripts. Collectively, our study deciphers the multi-layered, cooperative principles of RNA degradation and provides an epitranscriptomic perspective for understanding how RNA fate is encoded at the single-molecule resolution.

## Introduction

The dynamic turnover of RNA, a tightly regulated balance between synthesis and degradation, is fundamental to establishing temporally and spatially resolved gene expression programs^1,2^. RNA degradation acts as a central regulatory hub, actively modulating mRNA levels in response to various signals to maintain expression homeostasis^3^. Dysregulation of RNA decay pathways has been implicated in diverse human diseases, including developmental disorders and cancer^4–6^. Therefore, dissecting the molecular mechanisms and regulators that involved in RNA degradation is essential for a comprehensive understanding of gene expression and its role in health and disease.

The precise regulation of mRNA degradation depends on a combination of *cis*-regulatory elements and molecular features encoded within each transcript^1,2,7^. The length of the 3′ poly(A) tail, dynamically regulated through deadenylation, often serves as the rate-limiting step for mRNA degradation^8,9^. At the 5′ end, the cap structure shields the transcript from exonuclease-mediated degradation, while its removal (often called decapping) serves as the trigger for subsequent 5′-to-3′ degradation^10^. In addition, internal sequence elements like AU-rich elements (AREs) in the 3′ UTR destabilize mRNA by recruiting specific RBPs^11,12^, and codon usage bias within the coding sequence can influence stability by modulating ribosome dynamics during translation^13^.

Recently, chemical modifications of RNA transcripts have emerged as a pivotal layer of post-transcriptional regulation^14–16^. Among the hundreds of known RNA modifications, N6-methyladenosine (m6A) is the most abundant and extensively studied internal modification in eukaryotic mRNA^16^. m6A affects the entire mRNA lifecycle, with one of its most well-established functions being the regulation of mRNA stability through promotion of mRNA degradation^17,18^. This degradation is typically mediated by reader proteins, such as the YTH family proteins, which recognize m6A and recruit downstream effector complexes, including the CCR4-NOT deadenylase and RNase P/MRP complexes, to accelerate mRNA decay^18–20^. In addition, recent studies have revealed that m6A within the coding sequence can also trigger mRNA degradation by inducing ribosome collisions during translation^21,22^.

Despite the established role of m6A in promoting RNA degradation, key questions persist regarding its context-dependent function: how its impact varies across genes, isoforms, and even specific transcript regions, and how it functionally integrates with other post-transcriptional layers to orchestrate RNA decay? Whether these features constitute an “epitranscriptomic code” that links a transcript’s intrinsic features to its fate is still an open question. Deciphering this multi-layered regulation has been constrained by conventional short-read sequencing (i.e., NGS), which captures regulatory features in a fragmented and sequence-population-averaged manner, thereby obscuring how multiple signals are integrated within individual RNA molecules. In contrast, nanopore direct RNA sequencing (DRS) offers a unique opportunity to overcome this limitation^23^. By directly sequencing full-length native RNA molecules, DRS simultaneously captures isoform structure, poly(A) tail length, and RNA modifications such as m6A within the same molecule^23,24^. This single-molecule, multi-dimensional resolution enables direct interrogation of how regulatory features are jointly organized on individual transcripts, providing a powerful framework to test whether RNA fate is governed by a multi-factorial epitranscriptomic code^25^.

In this study, we leverage nanopore DRS to interrogate RNA decay at single-molecule resolution during a time-resolved transcriptional inhibition assay. We find that m6A regulates RNA decay through a stoichiometry-dependent mechanism, in which modification level quantitatively shapes degradation kinetics and drives a selective depletion of highly methylated RNA molecules over time. At the isoform level, we identify Regional m6A Clusters (RMCs) as regulatory elements that link alternative splicing to differential isoform-specific RNA stability. Finally, we develop a machine learning model that reveals a hierarchical contribution of sequence, poly(A), and modification features in determining RNA decay rates. Together, these results support a unified model in which RNA fate is encoded through the stoichiometric and combinatorial organization of regulatory features at the single-molecule level.

## Results

### DRS Accurately Quantifies mRNA Degradation Dynamics

We established an experimental pipeline using nanopore direct RNA sequencing (DRS) to profile transcriptomic changes in HEK293T cells following transcriptional inhibition with Actinomycin D (ActD). Poly(A)+ mRNA was collected at 0, 3, and 6 hours post-treatment, with *Drosophila* S2 cell mRNA added as an external spike-in to normalize expression levels across time points (Fig. 1a). Sequencing was conducted using the latest RNA004 chemistry from ONT platform, yielding high-quality data with substantial throughput. The resulting long reads achieved N50 values ranging from 1,635 to 1,905 bp and a maximum length of 133 kb (Supplementary Table 1), covering most full-length transcripts.

**Figure 1.**
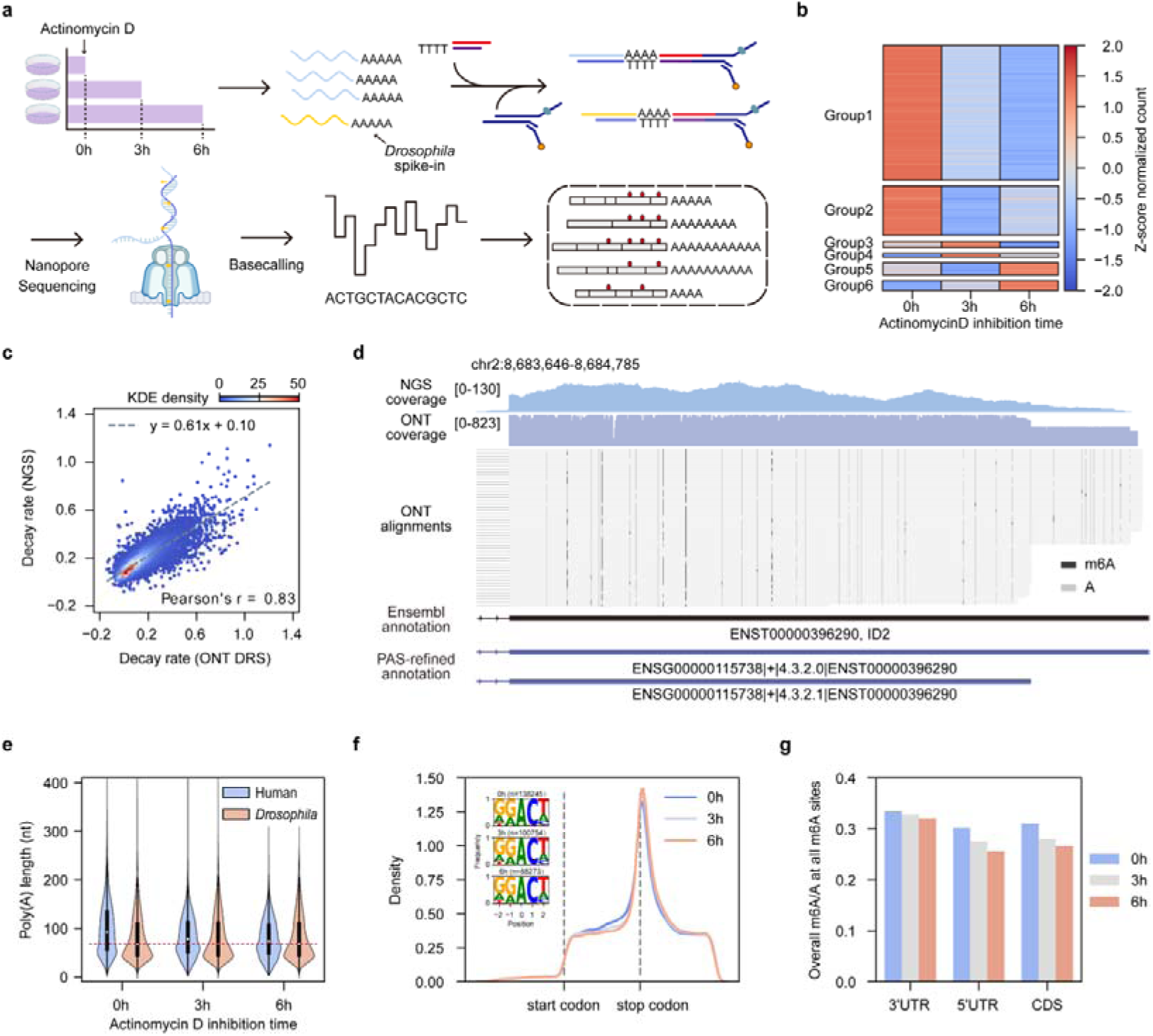
Nanopore Direct RNA Sequencing (DRS) enables comprehensive analysis of RNA degradation dynamics. **a**, Schematic of the experimental workflow. **b**, Heatmap of Z-score normalized expression for human genes, clustered into six groups (Groups 1-6) based on their temporal profiles after Actinomycin D treatment. **c**, Scatter plot comparing RNA decay rates calculated from nanopore DRS and NGS, demonstrating a strong correlation between the two methods (Pearson’s r = 0.83). **d**, Integrative Genomics Viewer (IGV) snapshot of the *ID2* locus, revealing at least two distinct polyadenylation sites (PASs) despite its annotation as a single isoform in Ensembl. Black and grey vertical bars represent m6A-modified and unmodified adenosines. **e**, Violin plots showing the progressive shortening of poly(A) tails in human transcripts (blue) over time, in contrast to the stable tails of *Drosophila* spike-ins (orange). The red dashed line marks the median poly(A) tail length of the spike-in. **f**, Metagene plot showing the distribution of m6A density across transcript regions at 0, 3, and 6 h. Dashed vertical lines indicate the start and stop codons. Insets show the sequence logos and the total number of identified m6A sites (*n*) for each time point. **g**, Bar plots representing the m6A/A ratios at all m6A sites within the 3′ UTR, 5′ UTR, and CDS at 0, 3, and 6 h after transcriptional inhibition.

Clustering analysis based on RNA expression dynamics revealed that the majority of genes (85.4%; groups 1 and 2) exhibited a marked decrease in abundance following transcriptional inhibition, consistent with active RNA degradation (Fig. 1b, Supplementary Fig. 1a, Supplementary Table 2). A smaller subset of genes (14.6%; groups 3-6) showed more stable expression (Supplementary Figs. 1b-d). Functional enrichment analysis revealed that these transcripts were predominantly involved in core biosynthetic processes, such as translation and respiration (Supplementary Fig. 1e), which is consistent with their exceptionally long half-lives. Critically, the RNA decay rates calculated from our DRS data were highly correlated (Pearson’s r = 0.83) with those derived from parallel short-read RNA-seq experiments (Fig. 1c), validating the accuracy of DRS in quantifying global RNA degradation dynamics.

### DRS Captures Multi-modal Transcriptome Features at Single-molecule Resolution

The long-read capability of DRS facilitates precise resolution of transcript architecture. Our analysis revealed that existing gene annotations often underestimate the complexity of alternative polyadenylation (APA). For instance, the *ID2* gene, previously annotated as a single isoform, utilizes at least two distinct polyadenylation sites (PASs), which would likely be missed by NGS (Fig. 1d, Supplementary Figs. 1f-g). To refine existing gene models, we used transcript 3′ end information from DRS data to generate a “PAS-refined annotation”, adding 3,558 novel PASs to 2,333 distinct splice variants (Methods, Supplementary Fig. 2a, Supplementary Table 3). Using this refined annotation, we re-evaluated transcriptome diversity and found that 2,665 splice variants (12.09%) utilize multiple PASs. In contrast, the original annotation identified this feature in only 1.51% of variants (Supplementary Figs. 2b-c). This precise PAS annotation allowed us to investigate the regulatory implications of APA, such as the use of variable 3′ UTRs. Notably, it also revealed instances of internal PAS usage within the coding sequences (CDS) of genes such as *CDC6* and *CCDC107*, which can potentially generate truncated protein products (Supplementary Figs. 2d-e).

In addition to re-evaluating isoform annotation, DRS enables the direct measurement of poly(A) tail length at single-molecule resolution. Following transcriptional inhibition, endogenous HEK293T mRNAs exhibited a progressive shortening of their poly(A) tails over time, whereas the poly(A) tails of *Drosophila* spike-in mRNA remained stable (Fig. 1e, Supplementary Figs. 3a-c), validating the ability of DRS to capture dynamic poly(A) tail remodeling during RNA decay. Based on these time-resolved measurements, we quantified the deadenylation rate for individual isoforms by modeling the temporal shortening of poly(A) tails (Methods). This provides a direct readout of RNA decay kinetics at isoform resolution, which is not accessible from steady-state measurements alone. Notably, we observed a positive correlation between poly(A) tail length and several isoform features, including transcript length, 3′ UTR length, number of exons, and mean exon length (Supplementary Figs. 3d-h). Previous studies have shown that longer transcripts containing more exons tend to require longer transcriptional elongation times and/or more complex splicing processes^26–28^. Such extended processing may result in prolonged nuclear retention^29^, which could provide more time for the addition of longer poly(A) tails.

A distinct advantage of DRS is its ability to directly detect RNA modifications from raw electrical signals. Our m6A calling results showed high concordance with the GLORI method^30^, gold standard for m6A detection at single-nucleotide resolution, confirming the reliability of site-level m6A identification in our dataset (Methods, Supplementary Fig. 3i). Across different time points of transcriptional inhibition, we identified 88,273-138,245 high-confidence m6A sites (methylation rate > 0.1), which were predominantly enriched within the canonical DRACH motif and clustered near the stop codon (Fig. 1f). Comparing m6A modification rates before and after transcriptional inhibition, we observed a global decrease in m6A levels, with a more pronounced reduction in the 5′ UTR and CDS, while m6A levels in the 3′ UTR showed only a marginal decline (Fig. 1g). Therefore, based on the long-read capability of DRS, we obtained a comprehensive dataset encompassing transcript architecture, RNA modifications, and poly(A) tail dynamics on single-molecule. This framework provides an opportunity to investigate how different molecular features jointly influence RNA fate.

### Absolute Quantification Reveals Stoichiometry-Dependent m6A-Mediated RNA Decay

Previous methods for identifying m6A predominantly relied on short-read sequencing, which captures sequence-population-averaged modification signals at specific sites or peak regions (Fig. 2a). However, dynamic processes such as RNA degradation and translation operate on full-length RNA molecules rather than site-level features. Therefore, we defined two new metrics to systematically quantify single-molecule stoichiometric feature: Modifications Per Molecule (MPM), representing the total number of m6A sites on an individual RNA molecule, and Modifications Per Kilobase per Molecule (MPKM), reflecting the length-normalized modification density (Figs. 2a-b).

**Figure 2.**
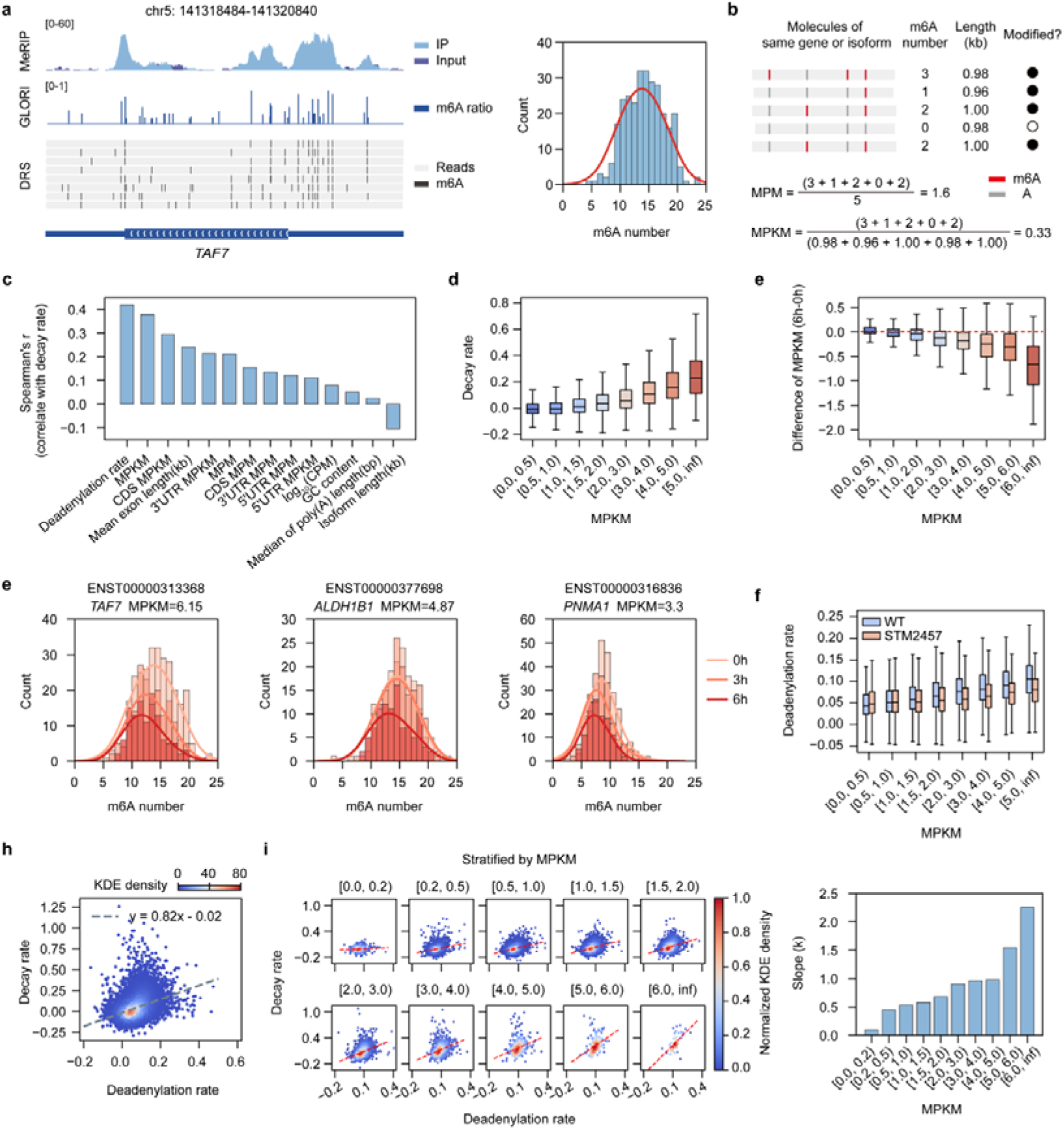
Single-molecule m6A stoichiometry is coupled with deadenylation and RNA decay. **a,** Comparison of m6A detection methods on the *TAF7* gene. Left, genome browser view show MeRIP-seq peaks (top), GLORI modification rates (middle), and individual DRS reads with identified m6A sites (black ticks, bottom). Right, distribution of m6A counts per molecule for *TAF7*. **b,** Schematic illustration of m6A quantification metrics. MPM (Modifications Per Molecule) is the average number of m6A sites per transcript. MPKM (Modifications Per Kilobase per Molecule) is the MPM normalized by transcript length. Filled and open circles indicate modified and unmodified transcripts, respectively. Red and grey vertical bars represent m6A-modified and unmodified adenosines. **c,** Spearman correlation coefficients between RNA decay rate and transcript features. **d,** RNA decay rates for isoforms grouped by m6A density (MPKM). **e,** Difference of m6A density (MPKM at 6 h minus 0 h) following transcriptional inhibition, stratified by initial MPKM. **f,** Distribution of m6A counts per molecule for three example genes (*TAF7*, *ALDH1B1*, and *PNMA1*) at 0h, 3h, and 6h after transcriptional inhibition. **g,** Deadenylation rates in wild-type (WT) and STM2457-treated cells, shown across isoform groups binned by MPKM. **h,** Scatter plot of RNA decay rate versus deadenylation rate. The dashed line represents a linear regression fit. **i,** Relationship between deadenylation rate and RNA decay rate stratified by MPKM. Left, scatter plots with robust linear regression using Tukey’s biweight function (red dashed lines). Right, slope (k) of the regression line for each MPKM bin. For all box plots, the center line denotes the median, box edges represent the interquartile range (IQR), and whiskers extend to 1.5× the IQR.

RNA decay is governed by a complex interplay of *cis*-regulatory elements and epigenetic modifications^1,2,7^. Conventional short-read approaches typically require multiple independent assays to capture these features, making it difficult to directly compare their relative contributions within a specific transcript. By contrast, DRS enables simultaneous acquisition of multi-modal transcriptomic features from the same molecules, including sequence composition, transcript architecture, absolute m6A abundance (MPM and MPKM), poly(A) tail length, and deadenylation rates. Integrating these features (Supplementary Table 4), we systematically evaluated their relationships with RNA decay rates. Both m6A abundance (measured by MPM or MPKM) and deadenylation rates showed the strongest positive correlations with RNA decay, identifying them as major determinants of RNA stability (Fig. 2c, Supplementary Fig. 4, Supplementary Fig. 5a).

Although m6A is widespread, with mammalian transcripts harboring an average of 3-5 sites^31,32^, whether individual sites act independently or cooperatively remains unclear. While some studies suggest that a single m6A site is sufficient for regulatory function^33,34^, others propose a stoichiometry-dependent model in which increasing numbers of m6A sites promote RNA decay^35–37^, underscoring the limitations of site-level analyses. To address this, we examined the relationship between m6A abundance and decay at the isoform level. We found that m6A abundance, quantified by either MPM or MPKM, positively correlated with RNA decay rates (Fig. 2d, Supplementary Fig. 5b), with highly modified isoforms exhibiting faster degradation kinetics (Supplementary Fig. 5c).

To validate this finding, we treated cells with the METTL3 inhibitor STM2457 and tracked RNA decay dynamics (Supplementary Table 1). Following inhibition, global m6A abundance was reduced to approximately 30% of control levels at the single-molecule level (Supplementary Fig. 5d). Correspondingly, the decay rates of highly modified transcripts were significantly reduced (Supplementary Fig. 5e). Consistent with this, during transcriptional inhibition, we observed a marked decrease in the average m6A level in the remaining RNA molecules of high-m6A isoforms, whereas low-m6A isoforms showed little change (Fig. 2e). This pattern indicates that RNA molecules with higher m6A abundance are preferentially degraded over time, supporting a stoichiometry-dependent role for m6A in RNA decay.

To further elucidate how the reduction in m6A levels arises during transcriptional inhibition, we examined the distribution of m6A across individual RNA molecules within each isoform. We found m6A sites are broadly distributed across the majority of molecules rather than enriched in small subsets of reads (Fig. 2a, right panel). Building on this baseline distribution, we next analyzed temporal changes in these molecular populations. In highly modified isoforms, the proportion of heavily modified RNA molecules progressively decreased during transcriptional inhibition (Fig. 2f, Supplementary Fig. 6). Importantly, because comparisons are made within isoforms, this analysis controls for sequence and structural variation. These results demonstrate that RNA molecules carrying higher numbers of m6A modifications are preferentially degraded, providing direct evidence that m6A-mediated RNA decay operates in a stoichiometry-dependent manner at the single-molecule level.

Notably, modification density (MPKM) exhibited a stronger correlation with decay rates than absolute modification counts (MPM), suggesting that local modification density is a more informative determinant of RNA stability (Supplementary Fig. 7a). This result highlights the importance of regional m6A distribution. Consistent with this, when examining different transcript regions, we found that m6A density within the coding sequence (CDS) was more strongly associated with decay rates than m6A in untranslated regions (UTRs) (Supplementary Fig. 7b). This observation is consistent with recently proposed translation-dependent decay models, in which ribosome stalling at m6A sites within the CDS serves as a key trigger for mRNA degradation^21,22,38^.

### m6A Stoichiometry Couples Deadenylation with RNA Decay

Deadenylation of the poly(A) tail is a critical initiating step in RNA degradation and often represents a rate-limiting process^8,9^. While previous studies have shown that m6A can promote deadenylation by recruiting the CCR4-NOT complex via the reader protein YTHDF2^19^, direct transcriptome-wide evidence linking m6A abundance to deadenylation rates remains limited. Our results revealed a significant positive correlation between m6A abundance and the rate of poly(A) tail shortening (i.e., deadenylation rate) at both the isoform and single-molecule levels (Fig. 2g, WT; Supplementary Figs. 8a-b). Notably, in contrast to previous observations^19,39^, inhibition of METTL3 did not significantly alter the global steady-state distribution of poly(A) tail lengths (Supplementary Fig. 8c). Instead, it led to a marked reduction in deadenylation rates, with a stronger effect observed in isoforms containing higher m6A levels (Fig. 2g, STM2457). These findings demonstrate that m6A primarily regulates the dynamic rate of poly(A) shortening rather than determining steady-state tail length, in a stoichiometry-dependent manner.

Consistent with previous studies, we observed a strong positive correlation between deadenylation rates and overall RNA decay rates (Fig. 2h). However, the strength of this relationship varied across transcripts. When isoforms were stratified by m6A density, those with very low m6A levels (e.g., MPKM < 1) exhibited generally low decay rates and showed little correlation between deadenylation and decay (Fig. 2i). In contrast, as m6A density increased, the association between deadenylation rate and decay rate progressively strengthened, accompanied by a marked increase in the slope of the linear relationship, indicating tighter kinetic coupling (Fig. 2i, right panel). Consistent with this, grouping transcripts by similar deadenylation rates further revealed that higher m6A abundance is associated with faster decay, particularly among transcripts with elevated deadenylation activity (Supplementary Fig. 8d). Together, these analyses support a functional synergy between m6A modification and the deadenylation process.

### Regional m6A Clusters (RMCs) Mediate Isoform-Selective RNA Degradation

Given the stoichiometry-dependent role of m6A in RNA decay, we reasoned that pairwise differences in m6A abundance between isoform variants from the same gene (isoform pairs) may contribute to variation in their degradation rates. Consistent with this, while most isoform pairs showed only modest differences in both m6A levels and decay rates (Supplementary Figs. 9a-b), those with the greatest divergence in m6A (top 20%) exhibited larger differences in degradation (Fig. 3a). This observation prompted us to further investigate the role of m6A in regulating isoform-specific decay.

**Figure 3.**
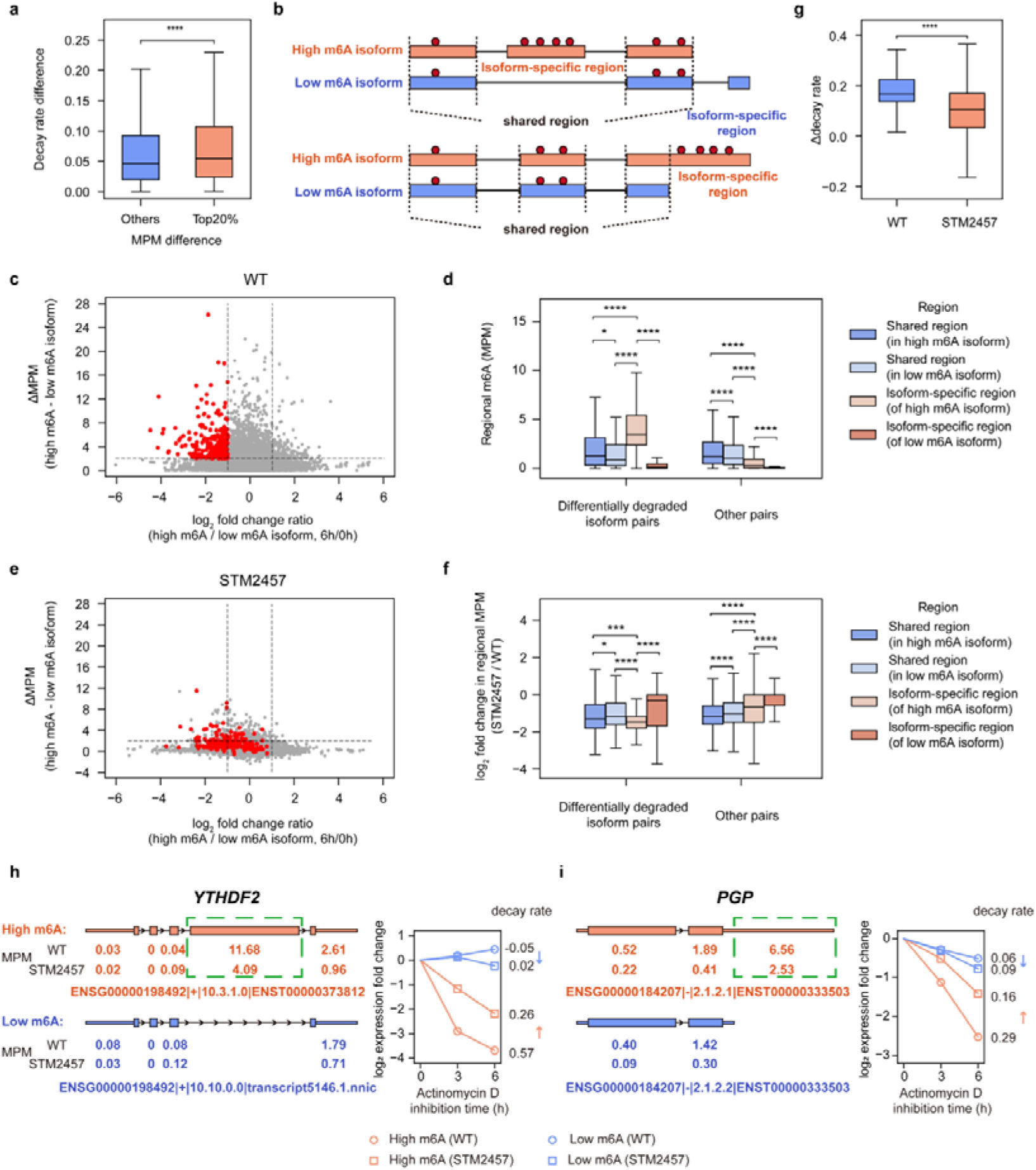
Region-specific m6A clusters (RMCs) regulate isoform usage preferences. **a,** Greater difference in decay rate for isoform pairs with the largest m6A (MPM) differences (Top 20%) compared to all other pairs (*\*\*\*\*P*<0.0001, Mann-Whitney U test). **b,** Schematic illustrating isoform pairs with shared and isoform-specific regions, marked with m6A sites (red dots). **c,** Volcano plot relating differential degradation (log_2_ fold change in abundance) to the difference in m6A levels (ΔMPM) for high-m6A versus low-m6A isoform pairs in wild-type (WT) cells. Red dots highlight pairs with faster degradation of the high-m6A isoform (log_2_FC < −1) and a large m6A difference (ΔMPM>2). **d,** Elevated m6A abundance (MPM) in isoform-specific regions compared to shared regions for the differentially degraded pairs identified in (**d**) (*\*P*<0.05, *\*\*\*\*P*<0.0001, Mann-Whitney U test). **e,** Volcano plot, similar to (**d**), but for cells treated with the METTL3 inhibitor STM2457. **f,** Box plots comparing the log_2_ fold change of regional m6A abundance (STM2457 vs. WT) in shared versus isoform-specific regions for the selected differentially degraded pairs versus all other pairs (*\*P*<0.05, *\*\*\*P*<0.001, *\*\*\*\*P*<0.0001, Mann-Whitney U test). **g,** Diminished difference in decay rates (Δdecay rate) between high-m6A and low-m6A isoforms in the selected pairs following STM2457 treatment (*\*\*\*\*P*<0.0001, Mann-Whitney U test). **h-i,** Examples of RMC-mediated regulation. **(h)** Alternative splicing of *YTHDF2* generates a stable, truncated isoform and an unstable, full-length isoform. **(i)** Alternative polyadenylation of *PGP* produces a stable, short-UTR isoform and an unstable, long-UTR isoform. For both genes, instability is linked to the presence of RMCs in the isoform-specific region (green dashed box) and is reversed by STM2457 treatment. For all box plots, the center line denotes the median, box edges represent the interquartile range (IQR), and whiskers extend to 1.5× the IQR.

To this end, we performed a pairwise analysis of isoforms from the same gene. For each gene, isoforms were classified into high-m6A and low-m6A counterparts based on steady-state MPM values (Fig. 3b). We then assessed whether differences in m6A levels between isoforms (ΔMPM) were associated with their relative changes in abundance following transcriptional inhibition, which reflects their differential degradation dynamics (Fig. 3c). This relationship was quantified as the log2 ratio of abundance changes between high- and low-m6A isoforms. While most of these pairs showed modest, a distinct subset exhibited both substantially higher m6A levels and faster degradation of the high-m6A isoform (ΔMPM > 2 and log2 fold change < −1; Fig. 3c, red dots). By aggregating data across all time points, we identified 356 such isoform pairs (from 223 genes) exhibiting this faster degradation trend for the high-m6A isoform (Supplementary Fig. 9c, Supplementary Table 5).

To determine the source of m6A differences in these pairs, we partitioned the exons of each pair into shared and isoform-specific regions and compared their m6A content (Fig. 3b). In the 356 differentially degraded pairs, the elevated m6A levels in the faster-degrading isoform were predominantly derived from isoform-specific regions. These regions showed significantly higher m6A levels than both the shared regions of the same transcript and the isoform-specific regions of the slower-degrading isoform (Fig. 3d). In contrast, m6A levels in shared regions were comparable between the two isoforms. These results indicate that localized enrichment of m6A within isoform-specific regions contributes to differential degradation. We therefore refer to these regions as Regional m6A Clusters (RMCs) and propose that they may function as a new regulatory element which mediating isoform-specific decay.

To further assess this model, we then re-examined the 356 differentially degraded isoform pairs in the context of METTL3 inhibition. After STM2457 treatment, differences in m6A levels between isoforms was markedly reduced, accompanied by a pronounced decrease in m6A within the RMC regions (Fig. 3e-f, Supplementary Fig. 9d). Intriguingly, the differential degradation between these isoform pairs was also significantly diminished upon METTL3 inhibition (Fig. 3g). Taken together, these results demonstrate that reducing m6A levels within RMCs attenuates their ability to mediate differential degradation, providing strong evidence that RMCs directly contribute to isoform-specific decay.

### RMCs Re-shape Isoform Expression Landscapes and Modulate Protein Coding Potential

Building on our finding that RMCs mediate differential isoform degradation, we next investigated the potential functional consequences of these differences at the level of protein output. Since isoform abundance directly influences the relative production of protein variants, RMC-mediated differences in decay rates may re-shape the protein-coding landscape. Analysis of the 356 isoform pairs revealed that in 264 cases, structural differences between isoforms resulted in altered protein products (Supplementary Figs. 9e-f), typically due to alternative splicing or APA events affecting the coding sequence. The remaining 92 pairs differed only in untranslated regions (UTRs).

The *YTHDF2* gene provides a representative example of how RMCs may influence protein output through alternative splicing (Fig. 3h, Supplementary Fig. 9e). The major isoform of *YTHDF2* includes exon 4, which encodes the YTH domain and is enriched for m6A, forming a typical RMC. A minor isoform lacking exon 4 produces a truncated protein without this domain. Under wild-type conditions, the major isoform is highly expressed but rapidly degraded, whereas the minor isoform is less abundant but more stable. Upon METTL3 inhibition with STM2457, the degradation of the major isoform was selectively slowed, while the minor isoform remained largely unchanged (Fig. 3h). Protein sequence analysis indicated that the exon-skipped isoform lacks most of the YTH domain and is therefore predicted to lose its m6A-binding capability (Supplementary Figs. 9g-h). These results suggest that RMC-associated m6A may influence the relative abundance of protein isoforms. Given the role of *YTHDF2* as an m6A reader, this differential stability may also represent a potential feedback mechanism in the regulation of m6A-mediated regulation.

Although RMCs located on UTR do not affect the protein-coding potential, they can regulate gene function by recruiting trans-regulatory factors such as miRNAs (Supplementary Figs. 9i-j). For example, the two major isoforms of *PGP* gene with distinct 3′ UTR lengths are generated via alternative PAS usage (Fig. 3i, Supplementary Fig. 9i). The longer isoform, derived from a distal PAS, contains an RMC and undergoes rapid degradation, whereas the shorter isoform lacks this feature and is more stable (Fig. 3i). Consistent with this pattern, METTL3 inhibition preferentially slowed the decay of the long-UTR isoform, supporting the idea that its instability is m6A-associated. Together, these examples illustrate that RMC-associated m6A can influence isoform abundance through multiple mechanisms, thereby re-shaping both protein-coding diversity and post-transcriptional regulatory potential.

### m6A Defines Distinct RNA Turnover Regimes without Directly Determining Expression Levels

Steady-state gene expression reflects the dynamic balance between RNA synthesis and degradation^40,41^. Consistent with previous studies, we observed only a weak negative correlation between m6A abundance and steady-state RNA expression across the transcriptome, with highly expressed isoforms generally exhibiting lower m6A levels and medium-to-low expression isoforms tending to carry higher m6A levels, albeit with substantial variability (Fig. 4a, Supplementary Fig. 10a).

**Figure 4.**
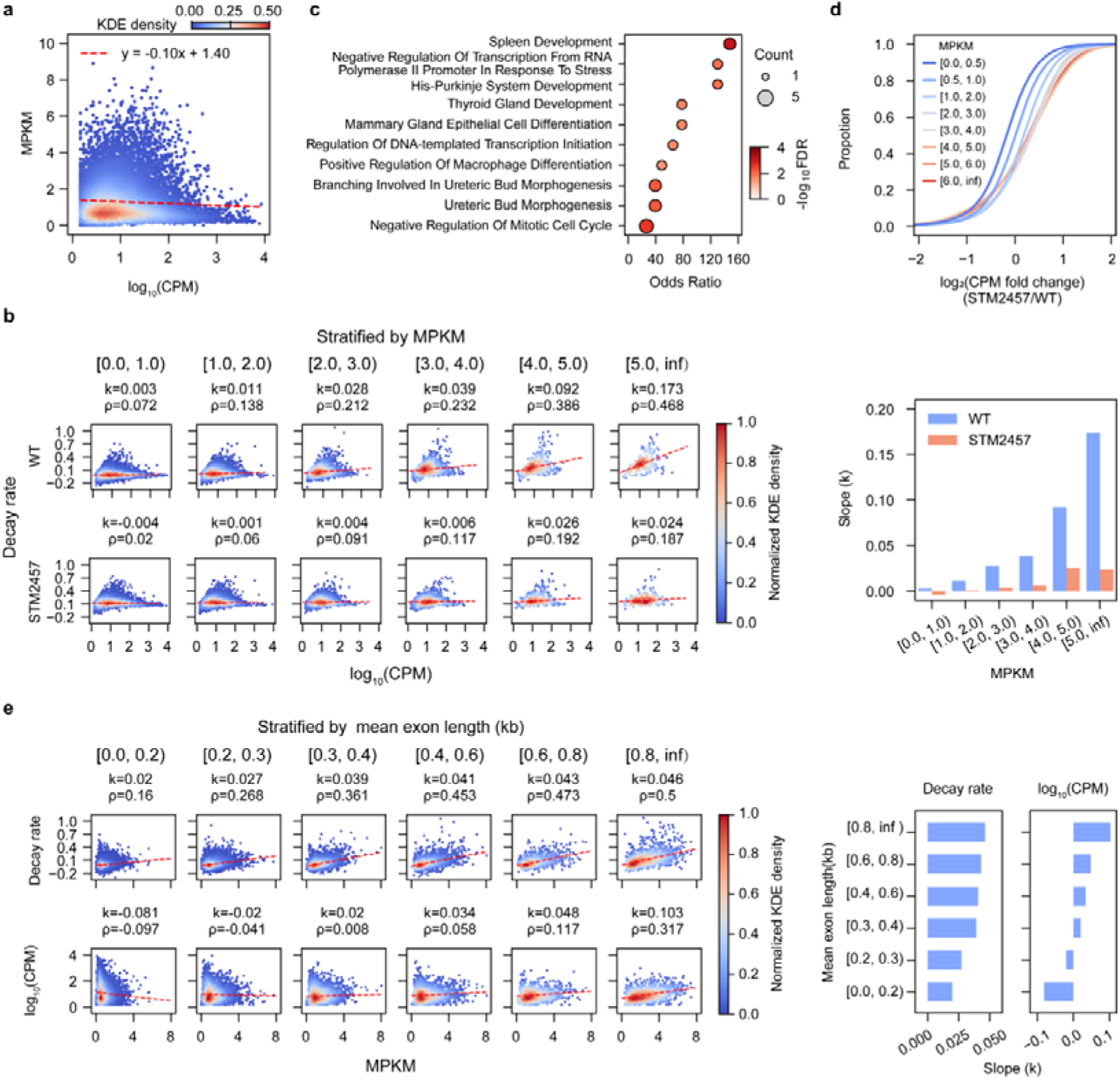
m6A density is associated with RNA turnover and transcript architecture. **a,** Scatter plot between m6A density (MPKM) and steady-state expression level (log_10_(CPM)) for all isoforms. **b,** Relationship between RNA decay rate and steady-state expression. Left, scatter plots for isoforms stratified by MPKM under wild-type (WT, top row) and STM2457-treated (bottom row) conditions. The slope of the regression line (k), derived from robust linear regression using Tukey’s biweight function, and the Spearman’s correlation coefficient (ρ) are indicated for each bin. Right, Bar plot of the regression slopes (k) for both conditions across the MPKM bins. **c,** Gene Ontology (GO) enrichment analysis for high-turnover isoforms (log_10_(CPM)>1, decay rate>0.4 and MPKM>3). Dot size corresponds to gene count, and color indicates statistical significance (-log_10_FDR). **d,** Cumulative distribution plots of log_2_(CPM fold change) after STM2457 treatment relative to WT, shown for isoforms stratified by their MPKM in the WT condition. **e,** Relationship of m6A density (MPKM) with decay rate and steady-state expression, stratified by mean exon length. Left, scatter plots showing MPKM versus decay rate (top row) and MPKM versus steady-state expression (log_10_(CPM), bottom row) across exon length bins. Values for k and ρ are defined as in (**b**). Right, bar plots of the regression slopes (k).

To further dissect this relationship, we stratified isoforms by m6A density and examined their kinetic properties. This analysis revealed that transcripts with different m6A levels adopt distinct RNA turnover regimes (Fig. 4b). Isoforms with very low m6A density (e.g., MPKM<1.0) displayed uniformly low decay rates, and their steady-state expression levels were largely uncoupled from decay rates, resulting in a broad dynamic range that includes highly expressed transcripts. In contrast, increasing m6A density was associated with progressively higher decay rates, along with a strengthening relationship between decay and steady-state expression (Fig. 4b, right panel). As a result, steady-state expression became increasingly constrained by degradation kinetics at higher m6A levels. Notably, within the high-m6A group, expression levels showed a clear positive correlation with decay rates, and a subset of transcripts exhibited both moderate-to-high expression and rapid decay. These observations indicate that high m6A modification does not simply correspond to low expression, but instead defines a class of transcripts characterized by rapid turnover.

Previous studies have shown that m6A is often enriched in transcripts requiring rapid and dynamic regulation, facilitating their timely turnover following activation^42,43^. In our dataset, we identified a subset of high-turnover transcripts within the high-m6A group, characterized by both elevated expression and rapid decay. Functional enrichment analysis revealed that these transcripts are associated with processes such as organ development, gene expression regulation, and cell cycle control (Fig. 4c). Consistently, canonical early response genes (ERGs) in our dataset also exhibited high m6A levels and rapid decay kinetics (Supplementary Fig. 10b). Upon global reduction of m6A levels using the METTL3 inhibitor STM2457, highly modified transcripts showed a pronounced decrease in decay rates accompanied by an increase in steady-state expression, whereas low-m6A transcripts were largely unaffected (Fig. 4d). These results indicate that high-turnover transcripts are particularly sensitive to m6A-dependent regulation, supporting a role for m6A in enabling rapid and reversible gene expression programs.

Recent studies have shown that long internal exons and specific splicing features influence m6A deposition, a pattern that is recapitulated in our DRS data^44–46^ (Supplementary Figs. 10c-d). Integrating these findings with our multi-modal DRS data, we linked transcript structure to downstream RNA kinetic behavior (Fig. 4e). Transcripts with longer exons were more likely to exhibit higher m6A density and were preferentially associated with a rapid turnover regime. These results suggest that RNA degradation kinetics may be partially encoded in gene architecture and implemented through m6A-mediated regulation.

### Machine Learning Reveals Key Determinants and Heterogeneity of RNA Degradation Control

To systematically evaluate the multi-layered determinants of RNA decay, we employed a machine learning framework. We curated a comprehensive feature set (101 in total) for each isoform, encompassing sequence composition (e.g., codon and k-mer frequencies), transcript structure, m6A modification, poly(A) tail dynamics, and expression level. A Gradient Boosting Tree model was trained to predict RNA degradation rates and quantify the relative contribution of these molecular features (Fig. 5a, Supplementary Table 6). The optimized model successfully captured variation in RNA degradation rates in held-out test set (Fig. 5b, Supplementary Fig. 11a), supporting the predictive value of the selected features and the multifactorial nature of RNA decay. Global feature importance analysis identified deadenylation rate, steady-state expression (log_10_(CPM)), and m6A density (MPKM) as the top predictors (Fig. 5c, Supplementary Fig. 11b, Supplementary Table 7), providing independent support for our earlier observations. Notably, m6A density within the coding sequence (CDS) emerged as a stronger predictor than its density in the 3′ UTR, a trend consistent with our correlation analyses and previous reports^21,22,38^.

**Figure 5.**
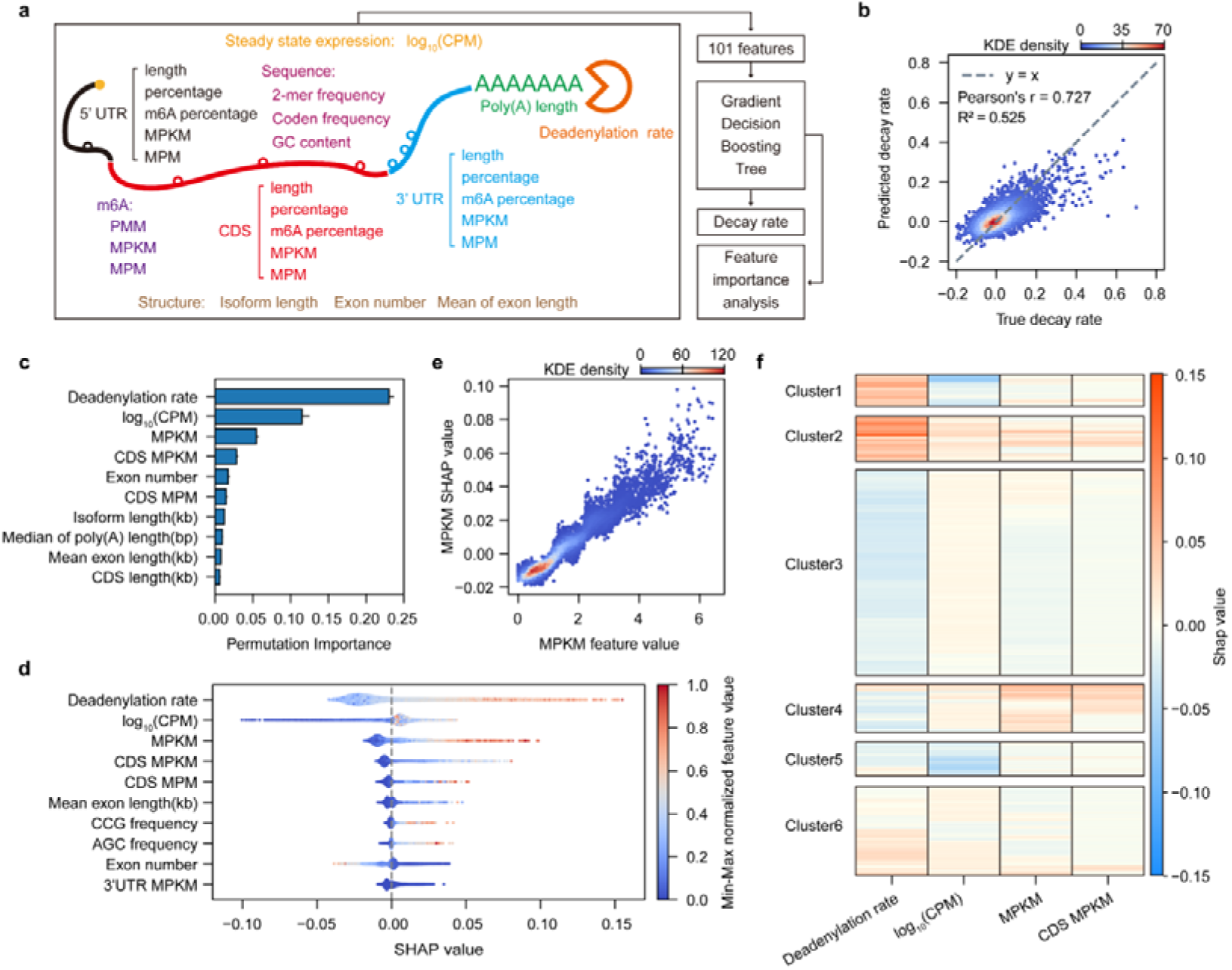
A machine learning model for the prediction and interpretation of multi-modal regulation in RNA degradation. **a,** Schematic of the overall workflow, encompassing feature extraction from DRS data, training of a Gradient Boosting Tree model, and model interpretation via feature importance and SHAP analysis. **b,** Comparison between model-predicted and experimentally determined decay rates for an independent test set. **c,** Top 10 most important features ranked by permutation importance on the test set. Error bars indicate the standard deviation from multiple permutations. **d,** SHAP summary plot for the top 10 features. Each dot represents an isoform with its x-position indicating the SHAP value (impact on model output) and its color representing the feature’s value (high or low). **e,** SHAP dependence plot for the MPKM feature, illustrating the relationship between a feature’s value (x-axis) and its corresponding SHAP value (y-axis) for each isoform. **f,** Hierarchical clustering of transcripts into six groups based on SHAP values for the four most influential features (Deadenylation rate, log_10_ (CPM), MPKM, and CDS MPKM).

While global feature importance provides an overall ranking, the contribution of these features can vary substantially across transcripts. To capture this heterogeneity, we performed SHAP (SHapley Additive exPlanation) analysis to quantify feature contributions at the individual isoform level^47^. Consistent with the global results, deadenylation rate, steady-state expression, and m6A density (both global and CDS-specific) remained dominant contributors (Fig. 5d). SHAP dependence plots, which show how the value of a feature impacts its contribution to the model’s prediction, allowed for a deeper analysis of these relationships. In the case of m6A density, this revealed a consistent positive relationship between m6A density and predicted degradation rate, indicating a model-inferred stoichiometry-dependent trend (Fig. 5e, Supplementary Fig. 11c). In contrast, other key features (such as deadenylation rate) exhibited complex, non-linear relationships (Supplementary Figs. 11d-e). Notably, some features with low global importance, such as the usage frequency of specific codons (e.g., CCG, AGC), showed contributions within specific subsets of isoforms (Fig. 5d), underscoring the context-dependent nature of degradation regulation.

To further characterize this heterogeneity, we clustered isoforms based on the SHAP values of the four most influential features. This analysis partitioned the isoforms into six clusters with distinct feature contribution profiles (Fig. 5f). Gene Ontology (GO) enrichment analysis showed that these clusters were also functionally coherent. For example, cluster 1 was enriched for genes related to cellular stress response and core molecular process regulation, while cluster 3 was associated with cellular respiration and metabolism (Supplementary Fig. 11f). This result suggests that isoforms with distinct biological functions may be governed by distinct combinations of degradation-related features.

## Discussion

The regulation of RNA decay is governed by a complex and multi-modal feature integrating sequence, structural features, chemical modifications, and poly(A) tail dynamics, which can be collectively referred to as the “epitranscriptomic code”. Deciphering this code has remained challenging due to the fragmented nature of conventional short-read approaches, which measure these features independently and at molecule-population-level resolution. Here, by leveraging nanopore direct RNA sequencing (DRS), we establish an integrated framework that captures full-length isoforms, m6A modification, and poly(A) tail length simultaneously on individual RNA molecules. The introduction of stoichiometric metrics (MPM/MPKM), improved PAS annotation, and direct estimation of deadenylation rates together provides a unique chance for dissecting RNA fate at unprecedented resolution.

This single-molecule approach allowed us to redefine the role of m6A in RNA decay. Within this framework, we show that m6A regulates RNA decay through a stoichiometry-dependent kinetic mechanism. Rather than acting as a binary signal, m6A quantitatively modulates deadenylation rates at the single-molecule level, establishing a continuous relationship between modification load and decay kinetics. Importantly, RNA decay is accompanied by a progressive shift in molecular composition, whereby highly methylated RNA molecules are preferentially degraded over time. Consistent with this, m6A and deadenylation form a functionally coupled regulatory axis, linking m6A abundance to the efficiency of poly(A) tail shortening and overall RNA turnover. At the systems level, this stoichiometry-dependent regulation gives rise to a spectrum of RNA turnover behaviors. Highly methylated transcripts are not necessarily depleted at steady state; instead, they frequently exhibit high-turnover regimes, characterized by concurrent rapid synthesis and degradation. This suggests that m6A-mediated regulation is better understood as tuning RNA flux rather than simply reducing transcript abundance, enabling dynamic responsiveness while maintaining transcriptome stability.

A key question is how m6A stoichiometry itself is determined. Our results point to transcript architecture as an important upstream contributor. We identify Regional m6A Clusters (RMCs) as isoform-specific features that disproportionately influence decay rates, linking alternative splicing to RNA stability control. In parallel, the strong association between internal exon length and m6A density suggests that structural features of transcripts may bias the distribution of m6A deposition, thereby shaping their susceptibility to rapid turnover. Together, these findings suggest that transcript architecture functions as an upstream determinant of m6A distribution, thereby influencing RNA decay kinetics. While our data do not establish strict causality, they support a model in which structural features encode a regulatory propensity landscape that biases transcripts toward distinct turnover regimes. This view is consistent with previous proposals that aspects of mRNA fate may be pre-configured during RNA processing, rather than determined solely post-transcriptionally^48^.

Given the multi-dimensional nature of RNA regulation, our machine learning framework serves not only as a predictive tool but also as a means to uncover underlying regulatory principles. The model consistently identified key determinants such as deadenylation rate and m6A density. Notably, SHAP-based analyses revealed substantial variability in feature contributions across transcripts, indicating that the relative importance of regulatory factors is highly context-dependent. This suggests that RNA decay is not governed by a single dominant mechanism but instead emerges from diverse combinations of features acting in a transcript-specific manner.

Several limitations of this study should be noted. The poly(A)-selection-based DRS approach restricts our analysis to polyadenylated RNAs, excluding important classes such as histone mRNAs and other non-polyadenylated transcripts. In addition, the absence of 5′ cap information prevents integration of cap-dependent decay mechanisms into our kinetic framework. Future advances in long-read sequencing, including poly(A)-independent library preparation, 5′ cap detection, and metabolic labeling strategies, will enable a more complete characterization of RNA lifecycles. Integrating these approaches with single-cell and spatial transcriptomics, as well as more advanced computational models, will provide deeper insight into the regulatory logic governing RNA turnover^49–51^. Ultimately, such efforts will be essential for deciphering the complete regulatory code of the RNA lifecycle and understanding its role in development, health, and disease.

## Methods

### Cell Culture and Sample Preparation for Sequencing

#### Cell Culture

HEK293T cells were cultured in DMEM (Corning) supplemented with 10% (vol/vol) fetal bovine serum (FBS, Gibco) and maintained at 37°C with 5% CO_2_. *Drosophila melanogaster* S2 cells were cultured at 25°C in Schneider’s Drosophila Medium (Gibco) supplemented with 10% (v/v) FBS. All cell lines were regularly tested and certified negative for mycoplasma.

#### RNA Decay Assay and Library Preparation

For the RNA decay Assay, HEK293T cells were seeded in 10 cm dishes and pre-treated for 24 hours with either the METTL3 inhibitor STM2457 (TargetMol, #T9060; 9μM) or DMSO as a control. Transcription was subsequently inhibited by the addition of actinomycin D (5 μg/ml; Aladdin, #A113142). Cells were then harvested at 0h, 3h, and 6h. For each time point, 10^7^ cells were lysed in TRIzol reagent (Thermo Fisher Scientific, #15596026) supplemented with 25 ng of *Drosophila* S2 cell mRNA as an external spike-in for normalization. Total RNA was extracted following the manufacturer’s protocol.

#### High-throughput sequencing

For Direct RNA Sequencing (DRS), poly(A)+ mRNA was isolated from total RNA using the Dynabeads mRNA Purification kit (Thermo Fisher Scientific, 61006). Libraries were subsequently prepared with the SQK-RNA004 kit following the manufacturer’s protocol (Oxford Nanopore Technologies). For parallel Next-Generation Sequencing (NGS) analysis, libraries were constructed and subsequently sequenced on an Illumina NovaSeq 6000 platform.

### DRS Data Processing and Gene-Level Analysis

#### Basecalling, Quality Control, and Mapping

Direct RNA sequencing (DRS) reads were basecalled using dorado basecaller (v0.8.0) (https://github.com/nanoporetech/dorado) with the ‘hac’ model, enabling poly(A) estimation and a minimum quality score of 7 (’--estimate-poly-a, --min-qscore 7’). Basecall quality was assessed using dorado summary (v0.8.0) and NanoPlot^52^(v1.43.0). The resulting reads were then mapped to the human (GRCH38) and *Drosophila* (BDGP6) reference genomes using dorado aligner (v0.8.0). Any supplementary alignments were subsequently filtered out using Samtools^53^(v1.6).

#### Gene Expression Quantification and Normalization

Gene expression was quantified at the isoform level using IsoQuant^54^(v3.6.2) with default parameters. For subsequent gene-level analysis, genes with ≤5 mapped reads were excluded. Read counts for the remaining genes were then normalized across time points using the *Drosophila* spike-in data. Specifically, normalization factors for the 3h and 6h samples were derived from the ratio of *Drosophila*-mapped reads relative to the 0h sample.

#### Clustering and Functional Analysis of Gene Expression Profiles

Following normalization, human genes (protein-coding and lncRNA based on biotype annotations) were grouped into six clusters based on their temporal expression profiles. For visualization, the expression profile of each gene was Z-score normalized and presented as a heatmap generated with the seaborn package (v0.12.2). Functional enrichment analysis for Gene Ontology (GO) and KEGG pathways was performed using gseapy^55^(v1.1.4).

### Isoform Annotation Refinement and Reads Assignment

#### Construction of PAS-refined Annotation

To improve the accuracy of isoform-level analyses, the reference annotation was refined by incorporating polyadenylation site (PAS) information from IsoQuant (v3.6.2) outputs. First, 3′ ends from reads uniquely assigned to annotated isoforms (full splice or single exon match) were aggregated for each distinct splicing pattern. Proximal 3′ ends (within a ±50 bp window) were iteratively merged to define PAS peaks, represented by the position with maximum read support. These peaks were further consolidated if closer than 100 bp, and any peak supported by fewer than 30 reads was discarded. To account for potential 5′ end imprecision in DRS data, isoforms sharing identical splicing patterns but with differing 5′ or 3′ ends were collapsed based on IsoQuant’s “extended_annotation.gtf”, retaining only the isoform with the most upstream 5′ end. The identified PASs were then assigned to these collapsed isoforms. A PAS was designated as ‘novel’ if its position differed from the original annotation by >100 bp; otherwise, the original PAS was kept. Finally, all transcripts in this refined annotation were systematically renamed to the format gene_id|strand|A.B.C.D|transcript_id to encode isoform and PAS details (A: total number of distinct splice variants for this gene, B: Index of the specific splice variant (1-based), C: Number of distinct Polyadenylation Sites (PASs) associated with the splice variant indicated by B, D: Index of the specific PAS (0-based; 0 indicates the reference PAS).

#### Read Assignment to the Refined Annotation

Long reads were assigned to unique isoforms in the PAS-refined annotation using Trmap^56^(v0.12.6), which classifies read-to-isoform relationships by splicing chain matches. A custom algorithm was implemented to resolve ambiguities and ensure high-quality assignments. For multi-exon reads with an exact splice match (’=’ relationship), any ambiguity from alternative polyadenylation was resolved by selecting the isoform that minimized the number of unaligned bases at the ends. Such reads were retained for analysis if their alignment discrepancy was <40% of the isoform length. For single-exon reads, assignment was based on containment (’c’ or ‘k’ relationships), again selecting the best-matching isoform and requiring an alignment discrepancy of <20% of the isoform length.

### Analysis of RNA Decay Dynamics

#### Calculation of RNA Decay Rates

RNA decay rates (*k*) at both gene and isoform levels were estimated by fitting the normalized time-course expression data to a first-order exponential decay model: A_t_ = A_0_ * e^−kt 57^. In this model, A_t_ represents the RNA abundance at time *t*, A_0_ is the initial abundance at *t*=0, and *k* is the decay rate constant. Optimal *k* values were determined by minimizing the residual sum of squares between observed and modeled abundances using the L-BFGS-B algorithm. The fitting was performed in Python (v3.11.8) using the scipy.optimize.minimize function (SciPy v1.10.1), with an initial guess of 0 and bounds of [-2.5, 2.5] for *k*.

#### Comparison with NGS-derived Decay Rates

Sequenced reads were mapped using HISAT2^58^(v2.2.1), and gene-level counts were quantified with featureCounts^59^(v2.0.6). These counts were subsequently normalized using the *Drosophila* spike-in data. Decay rates were calculated from the normalized NGS data using the same exponential model and were compared with the DRS-derived rates.

#### Differential Isoform Decay and Functional Impact Analysis

To investigate differential decay within genes, isoforms were analyzed in a pairwise manner. First, the expression fold change (expFC) for each isoform *i* at time *t* relative to its steady-state level (*t*=0) was calculated: expFC_(t, isoform i)_= normalized counts_(t, isoform i)_/ normalized counts_(0, isoform i)_ Next, to assess m6A-dependent differential decay, isoform pairs were categorized as ‘high-m6A’ or ‘low-m6A’ based on their MPM values. The relative change in their expFC was then computed as a log_2_ ratio:

log_□_expFC ratio_(t, isoform pair)_ = log_□_(expFC_(t, high m6A isoform)_/ expFC_(t, low m6A isoform)_)

A negative value for this log_2_ ratio indicates a faster turnover of the high-m6A isoform relative to its low-m6A counterpart.

To understand the functional consequences of differential decay, the coding sequences (CDS) of these isoform pairs were compared. We established a comparison hierarchy utilizing either annotated CDS from our refined annotation or *de novo* predicted ORFs from orfipy^60^(v0.0.4). The primary comparison was between annotated CDSs if available for both isoforms. If not, predicted ORFs were compared against annotated CDSs, or against each other in a comprehensive pairwise manner if no annotations existed. An identical CDS/ORF match at any level of this hierarchy classified the isoform pair as encoding the same protein, allowing for an assessment of how differential degradation impacts the proteome.

### m6A Detection, Validation, and Quantification

#### m6A Calling

The detection of m6A sites from DRS reads was performed using Singlemod under user’s guidance^61^ (https://github.com/xieyy46/SingleMod-v1).

#### Isoform-Level Quantification

Three metrics were calculated to quantify m6A levels for each isoform: the Percentage of Modified Molecules (PMM), defined as the proportion of reads with at least one m6A call; the average number of m6A modifications per molecule (m6A Modifications Per Molecule, MPM); and the average number of m6A modifications per 1,000 base pairs (m6A Modifications Per Kilobase per Molecule, MPKM). These metrics were calculated using custom scripts.

### Poly(A) tail analysis

#### Estimation of Poly(A) Tail Length

The poly(A) tail length of each individual DRS read was estimated by the Dorado basecaller via the --estimate-poly-a parameter. For each isoform with ≥5 reads per sample, the median poly(A) tail length was calculated at each time point (0h, 3h, 6h).

#### Calculation of Deadenylation Rate

The rate of poly(A) tail shortening (deadenylation rate, λ) was estimated at the isoform level by fitting the median poly(A) tail lengths over the time course (0h, 3h, 6h) to a first-order exponential decay model: L_t_ = L_0_*e^−λt 57^. In the model, L_t_ is the median length at time *t*, L_0_ is the initial length at *t*=0, and λ is the deadenylation rate constant. Optimal λ values were determined by minimizing the residual sum of squares using the L-BFGS-B algorithm. The fitting was performed in Python (v3.11.8) using the scipy.optimize.minimize function (SciPy v1.10.1), with an initial guess of 0 and bounds of [-2.5, 2.5] for λ.

### Machine Learning Models for Decay Rate Prediction

#### Feature Engineering and Dataset Preparation

A set of 101 features were curated for each protein-coding transcript, encompassing sequence, structure, m6A modification, poly(A) tail, and expression level characteristics. To ensure data quality, this dataset was filtered to remove outliers based on predefined thresholds for key features (decay rate > −0.3 and ≤ 0.8; MPM ≤ 19; MPKM ≤ 6.5; deadenylation rate ≥ −0.1 and ≤ 0.3; median poly(A) length > 50 and ≤ 200 nt; and log_10_(CPM) ≤ 3.1; Supplementary Table 6). The final filtered dataset was then stratified by decay rates and randomly split into training (80%) and test (20%) sets using StratifiedShuffleSplit from scikit-learn (v1.2.2).

#### Model Training, Optimization, and Evaluation

A Gradient Boosting Regressor model was trained to predict RNA decay rates, chosen for its ability to capture non-linear relationships and feature interactions. Hyperparameter optimization was performed on the training set using a 5-fold cross-validated randomized search (sklearn v1.2.2, function: sklearn.model_selection.RandomizedSearchCV; arguments: n_iter=200, cv=5). The final optimized hyperparameters used for the model were: random_state=42, n_estimators=500, min_samples_split=48, min_samples_leaf=11, max_features=14, max_depth=9, and learning_rate=0.03. Model performance was evaluated on the held-out test set by calculating the Pearson correlation coefficient between predicted and actual decay rates.

#### Model Interpretation and Feature Importance

To interpret the model and identify key determinants of RNA decay, three complementary approaches were employed: (1) impurity-based feature importance from the fitted model; (2) permutation importance (sklearn v1.2.2, function: sklearn.inspection.permutation_importance, arguments: n_repeats= 10); and (3) SHapley Additive exPlanations (SHAP) values derived from cooperative game theory (shap v0.47.1, function: shap.TreeExplainer).

#### Clustering and Functional Analysis of Regulatory Patterns

To identify distinct regulatory patterns, transcripts were subjected to hierarchical clustering based on the SHAP values of the four most important features (deadenylation rate, log_10_(CPM), MPKM, and CDS MPKM). Clustering was performed using Ward’s linkage method (scipy v1.10.1, function: scipy.hierarchy.linkage), partitioning the transcripts into six clusters based on a distance criterion (scipy v1.10.1, function: scipy.hierarchy.fcluster; arguments: t=1.2, criterion=’distance’). Genes within each cluster were then analyzed for functional enrichment of Gene Ontology (GO) and KEGG pathways using gseapy (v1.1.4, function: gseapy.enrichr).

## Supplement information

Supplementary Figures 1-11

Supplementary Tables 1-7

## Data availability

Sequencing data generated in this work can be found in GEO with the accession number GSE301054.

## Acknowledgements

This work was supported by the National Key R&D Program of China (2022YFA0912900 and 2022YFC3400400), Guangdong Basic and Applied Basic Research Foundation (2024B1515020115), National Natural Science Foundation of China (32425034, 92253202, 32271499 and 32270644), Fundamental Research Funds for the Central Universities (24xkjc017, 23lgbj011), and Guangzhou Science and Technology Project (2024A04J3408).

## Author Contributions

G.-Z.L. and Z.Z. conceived the project; Z.Z. and C.-L.W. wrote the manuscript; Z.Z. and C.-L.W. analyzed the data with the assistance from Y.-Y.X., Z.-D.Z., G.-R.T., Z.-H.R., Z.-S.Q., T.-W.S., Y.-L.L., J.-T.H., Q.-Y.W. and J.-W.K.; Z.-Z.H, H.-Y.F. conducted the experiments with the assistance from H.-X.C., R.-J.L.,W.-Q.L. and F.W.; G.-Z.L. revised the manuscript. All authors reviewed the results and approved the final version of the manuscript.

## Competing Interests

The authors declare no competing financial interests.

## Notes

### Competing Interest Statement

The authors have declared no competing interest.

