## Supplementary Information for "Deciphering the epitranscriptomic code of RNA degradation with nanopore direct RNA sequencing"

### **Supplementary Figures**

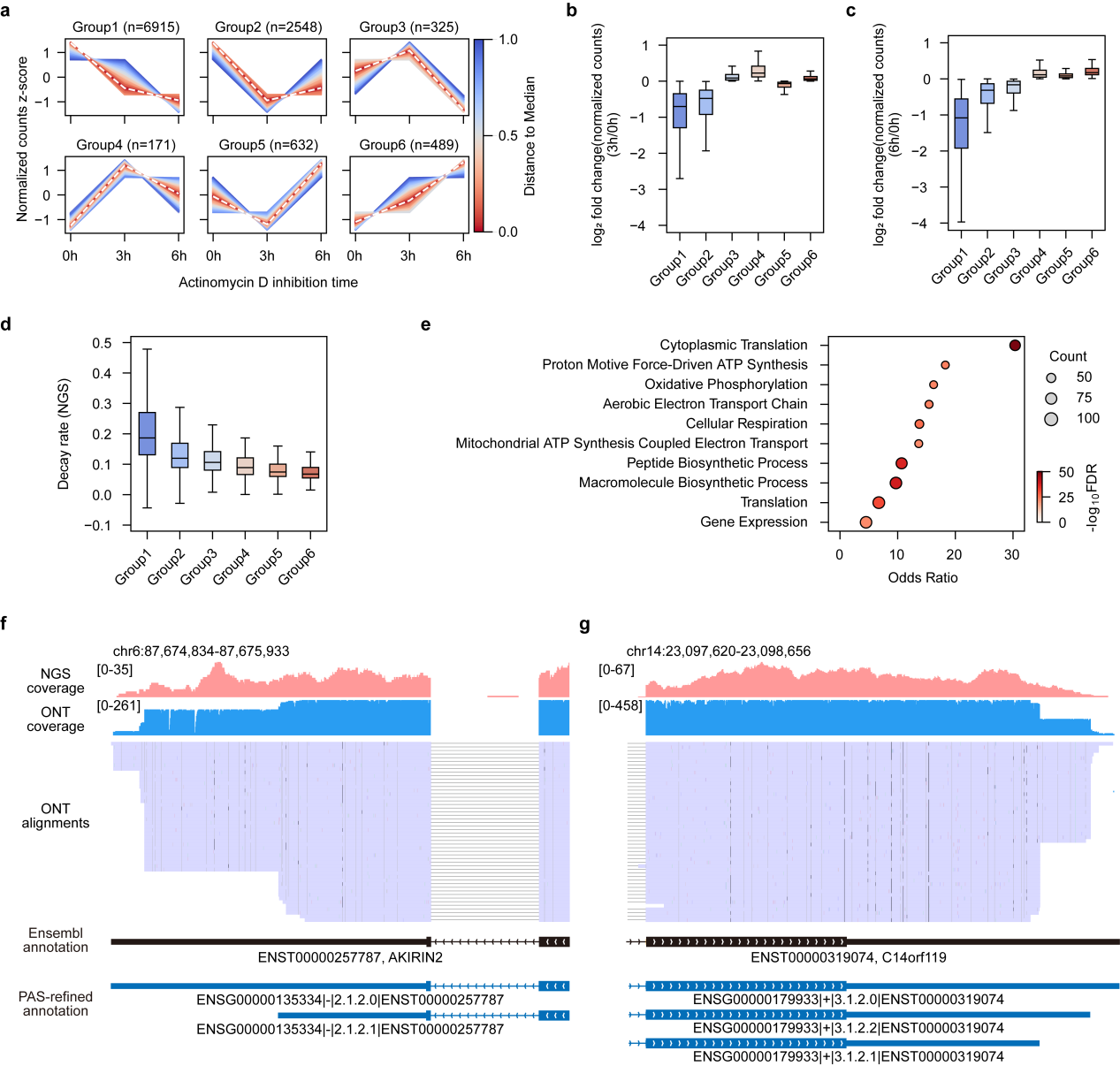
**Supplementary Fig. 1 | Characterization of global RNA degradation dynamics. a,** Temporal expression profiles of the six gene clusters from Fig. 1b following Actinomycin D treatment. Expression is Z-score normalized, with the number of genes in each group indicated. **b-c,** Log_2_ fold change of normalized transcript counts at 3h (**b**) and 6h (**c**) relative to 0h for each gene cluster. **d,** Distribution of RNA decay rates derived from parallel short-read RNA-seq experiments for the six gene clusters. **e,** Gene Ontology (GO) enrichment analysis of stable transcripts (Groups 3-6). Dot size is proportional to the number of genes (Count), and color intensity reflects the statistical significance (-log_10_FDR). **f-g,** Integrative Genomics Viewer (IGV) snapshots of the *AKIRIN2* (**f**) and *C14orf119* (**g**) loci reveal novel alternative polyadenylation (APA) sites. The tracks display read coverage from NGS and ONT, the Ensembl annotation, and our PAS-refined annotation.

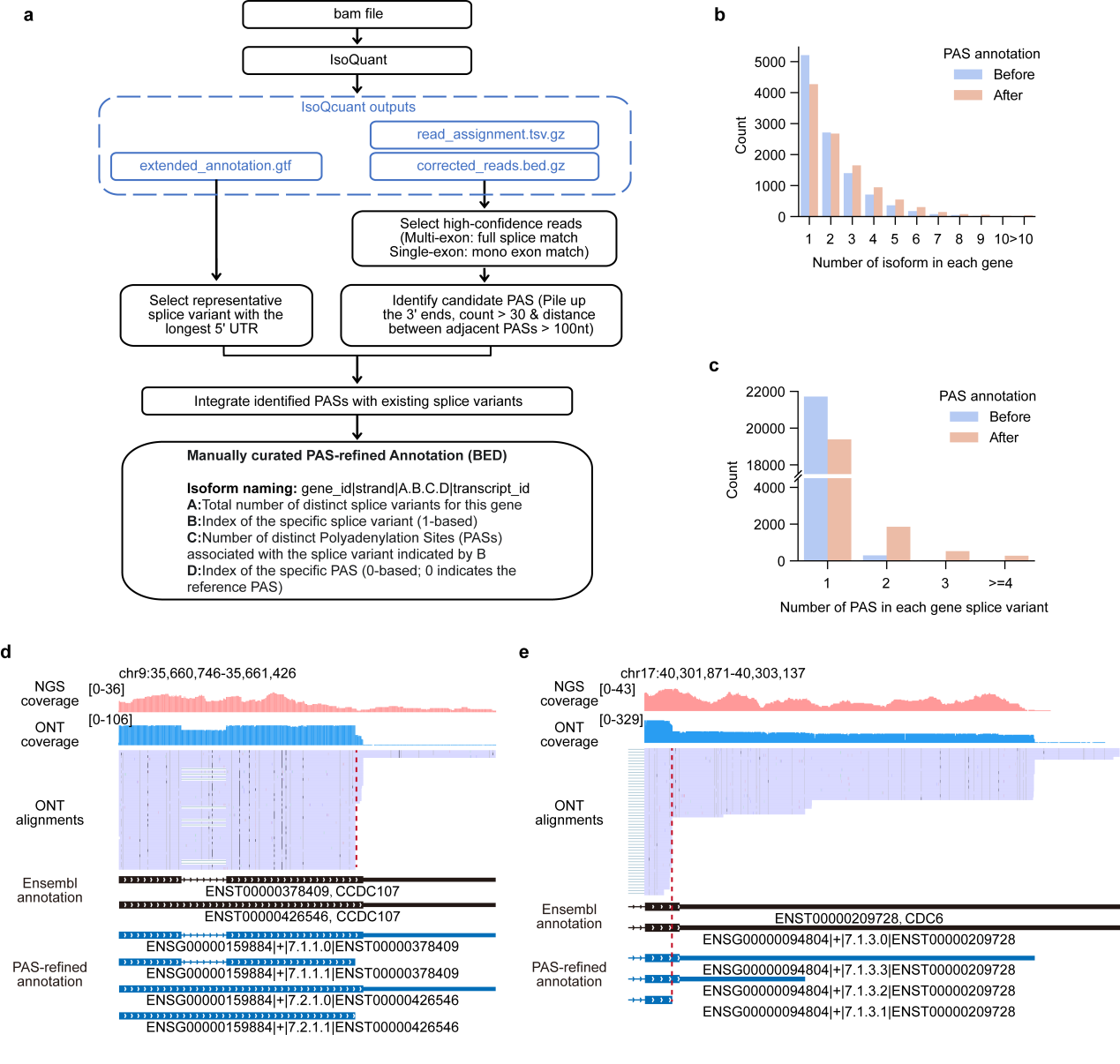
**Supplementary Fig. 2 | Construction of a PAS-refined annotation and analysis of isoform diversity. a**, Schematic of the pipeline for constructing the PAS-refined annotation, which integrates transcript 3′ end information from DRS data with existing splice variants. **b-c**, Comparison of the number of isoforms per gene (**b**) and distinct polyadenylation sites (PASs) per splice form (**c**) before and after annotation refinement. **d-e**, Refined annotation reveals extensive alternative polyadenylation, including internal PAS usage within the coding sequence (CDS) of genes such as *CCDC107* (**d**) and *CDC6* (**e**), shown in IGV snapshots.

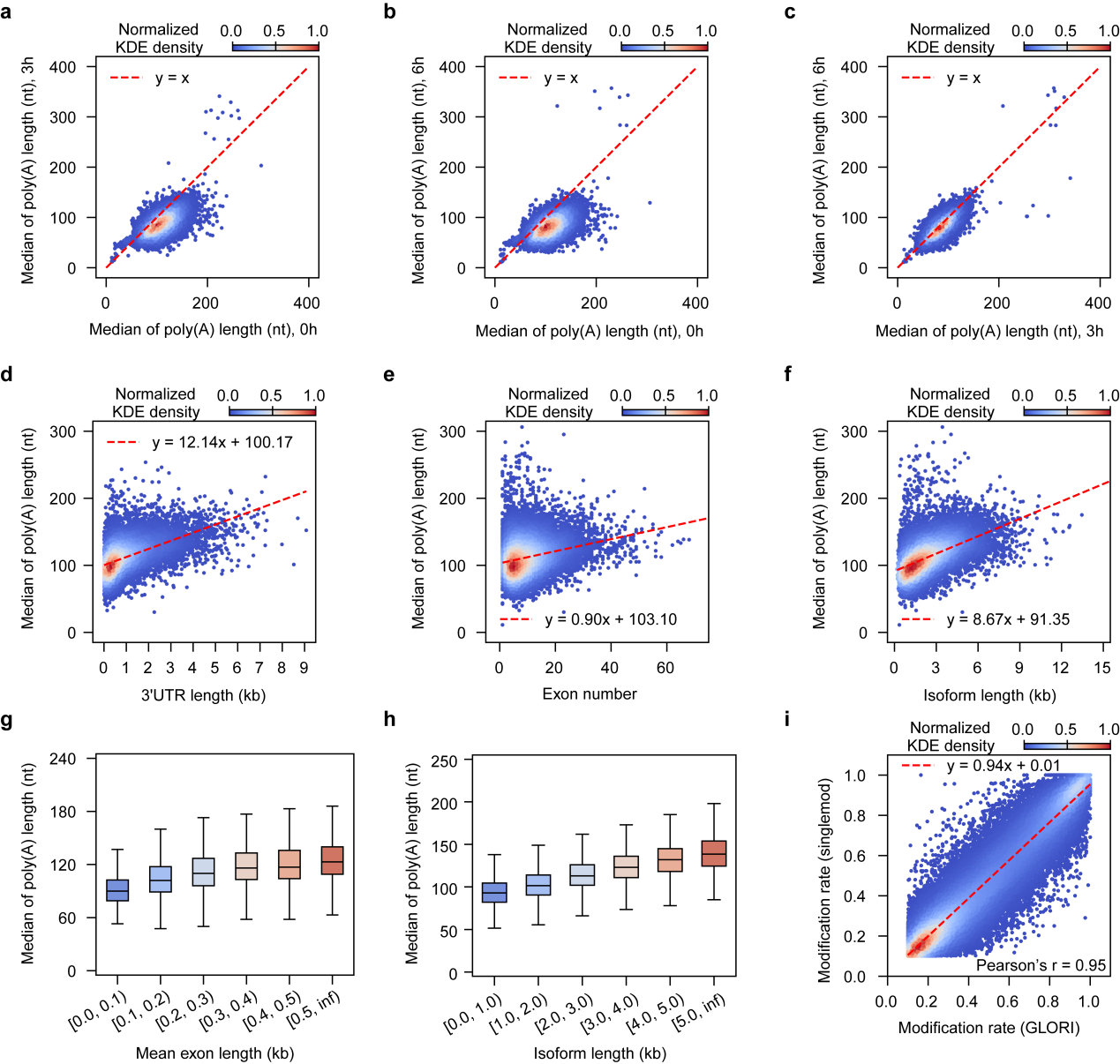
**Supplementary Fig. 3 | Characterization of poly(A) tail dynamics. a-c,** Comparison of median isoform poly(A) tail lengths between different time points: 3h vs 0h (**a**), 6h vs 0h (**b**), and 6h vs 3h (**c**). The red dashed line represents y=x. **d-f,** Scatter plots showing the relationship between median poly(A) tail length (at 0h) and 3′ UTR length (**d**), exon number (**e**), and total isoform length (**f**). **g-h,** Box plots of median poly(A) tail lengths for human isoforms, binned by mean exon length (**g**) and isoform length (**h**). **i**, Scatter plots comparing site-level m6A modification rates between ONT DRS and NGS-based GLORI method. The red dashed line represents a linear regression fit, and the Pearson's correlation coefficient is indicated. For all box plots, the center line denotes the median, box edges represent the interquartile range (IQR), and whiskers extend to 1.5× the IQR.

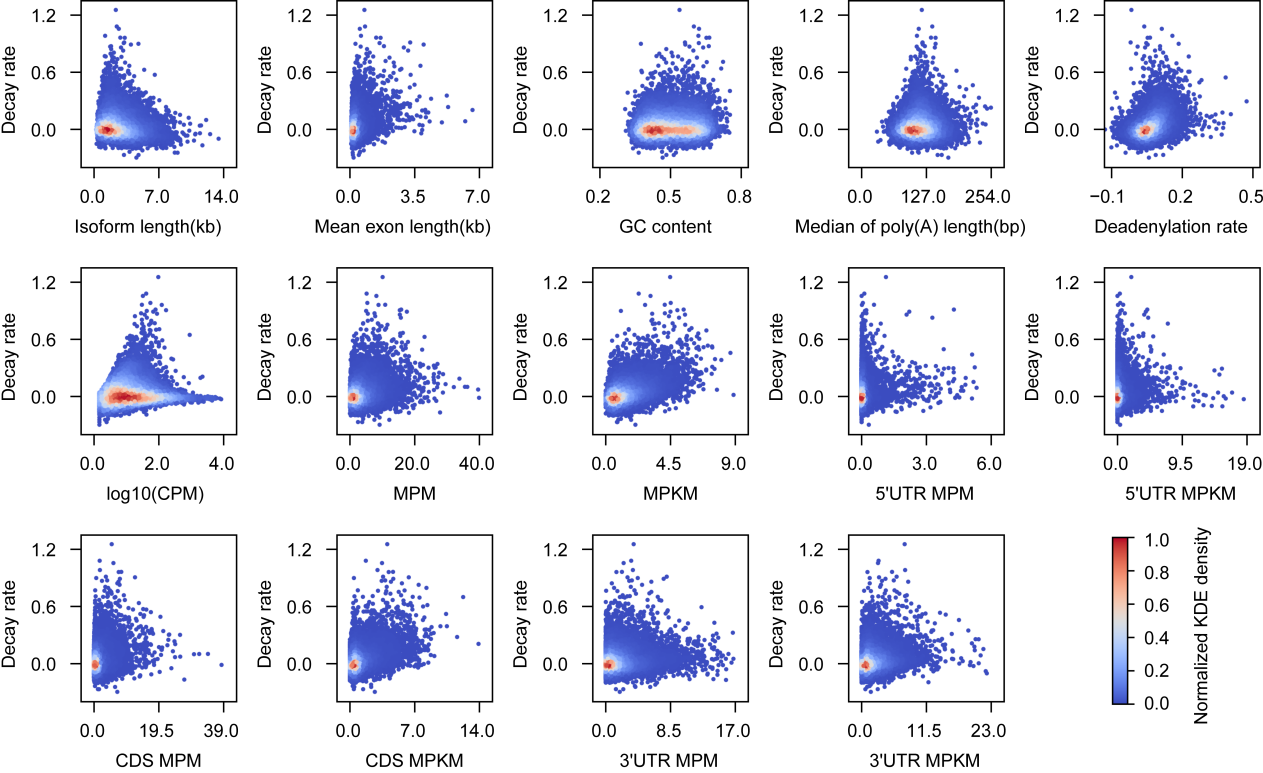
**Supplementary Fig. 4 | Relationship between RNA decay rate and various transcript features.** Scatter plots visualizing the relationship between RNA decay rate and 14 distinct transcript features. The features include structural properties (Isoform length, Mean exon length, GC content), steady-state expression (log_10_(CPM)), poly(A) tail dynamics (Median of poly(A) length, Deadenylation rate), and m6A stoichiometry (total MPM and MPKM, as well as region-specific metrics for 5′ UTR, CDS, and 3′ UTR). In each plot, the color intensity represents the normalized KDE density.

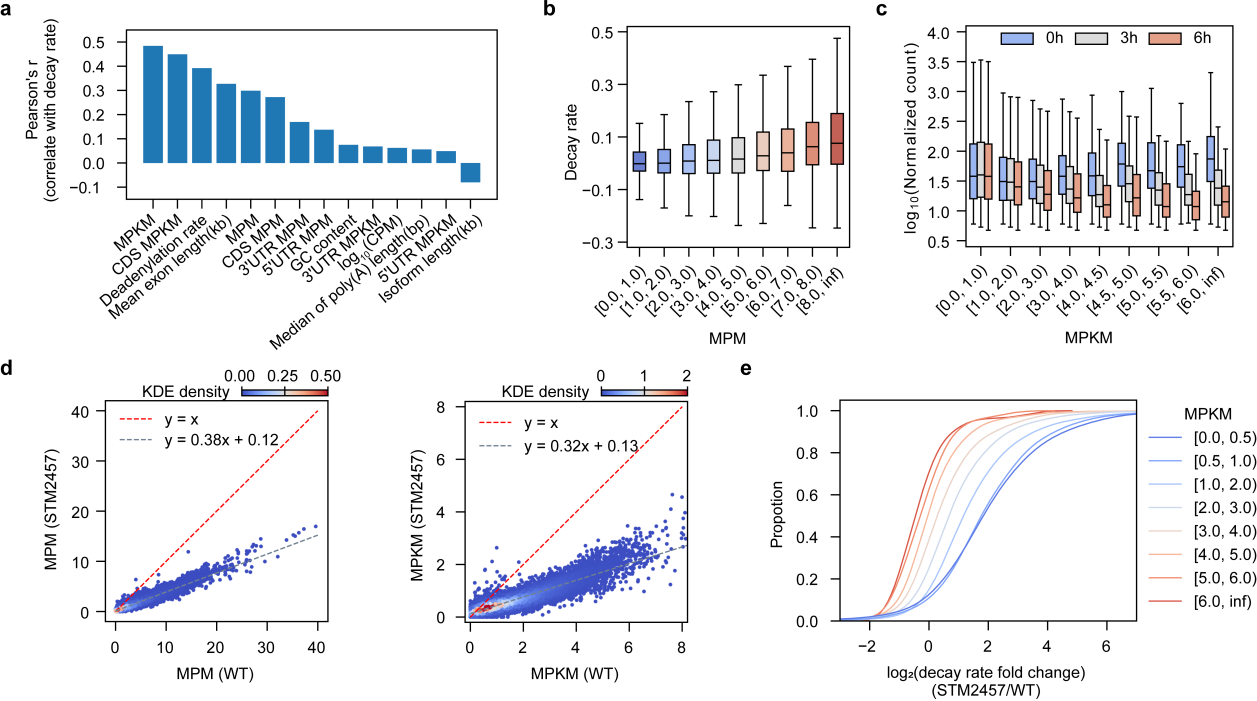
**Supplementary Fig. 5 | Relationship between m6A abundance and RNA decay dynamics. a,** Bar plot showing Pearson's correlation coefficients (r) between various transcript features and decay rate. **b,** RNA decay rates of isoforms stratified by m6A level (MPM). **c,** Box plots of log_10_ normalized transcript counts at 0h, 3h, and 6h after transcriptional inhibition, for isoforms binned by m6A density (MPKM). **d,** Comparison of MPM and MPKM between wild-type (WT) and STM2457-treated cells. The red dashed line indicates y=x, and the grey dashed line represents a linear regression fit. **e**, Cumulative distribution plots of the log_2_(decay rate fold change) between STM2457-treated and WT cells, shown for isoforms stratified by WT MPKM.

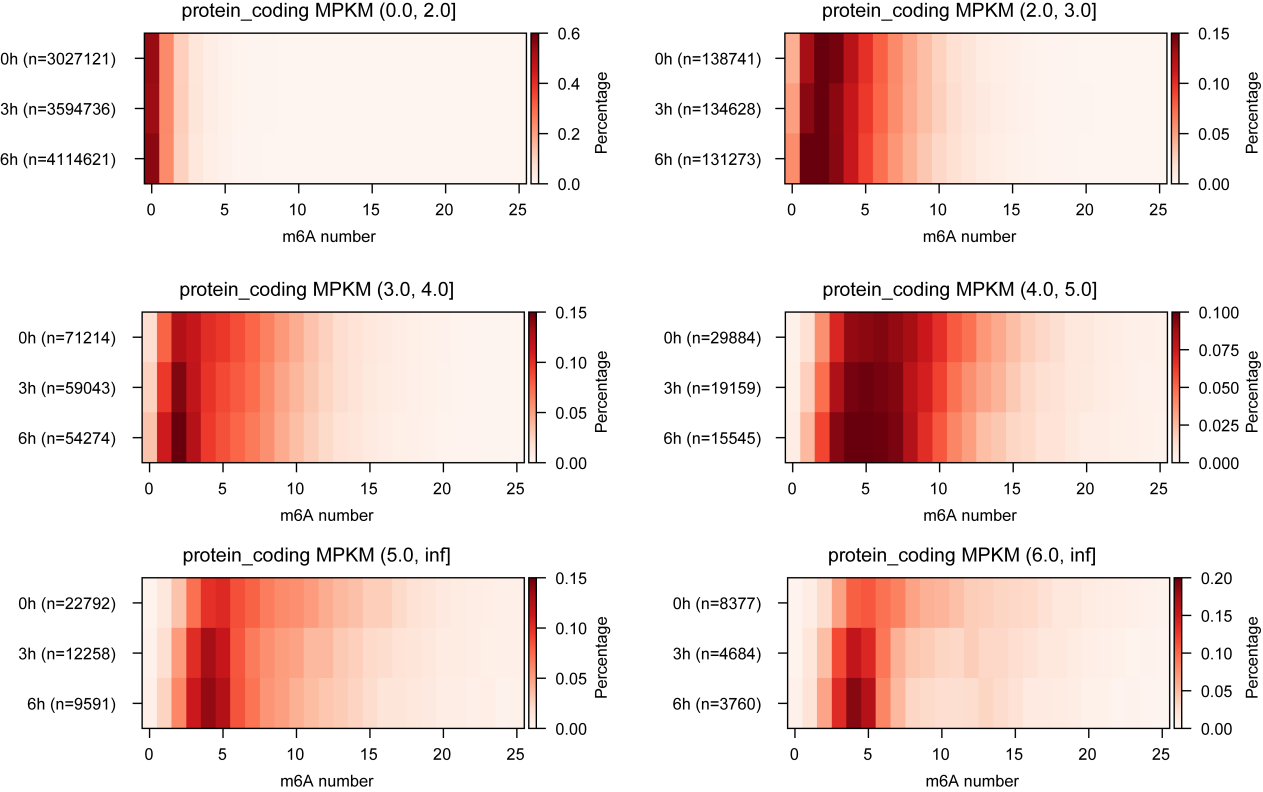

**Supplementary Fig. 6 |** **Dynamic shifts in single-molecule m6A stoichiometry during RNA decay.** Heatmaps illustrating the distribution of m6A modification counts per molecule at 0h, 3h, and 6h following transcriptional inhibition. Isoforms are stratified into six panels based on their modification density (MPKM). For each time point, the total number of analyzed molecules (*n*) is indicated. The color intensity represents the percentage of the molecular population containing a specific number of m6A sites (x-axis), showing a progressive depletion of heavily modified molecules over time in the high-MPKM categories.

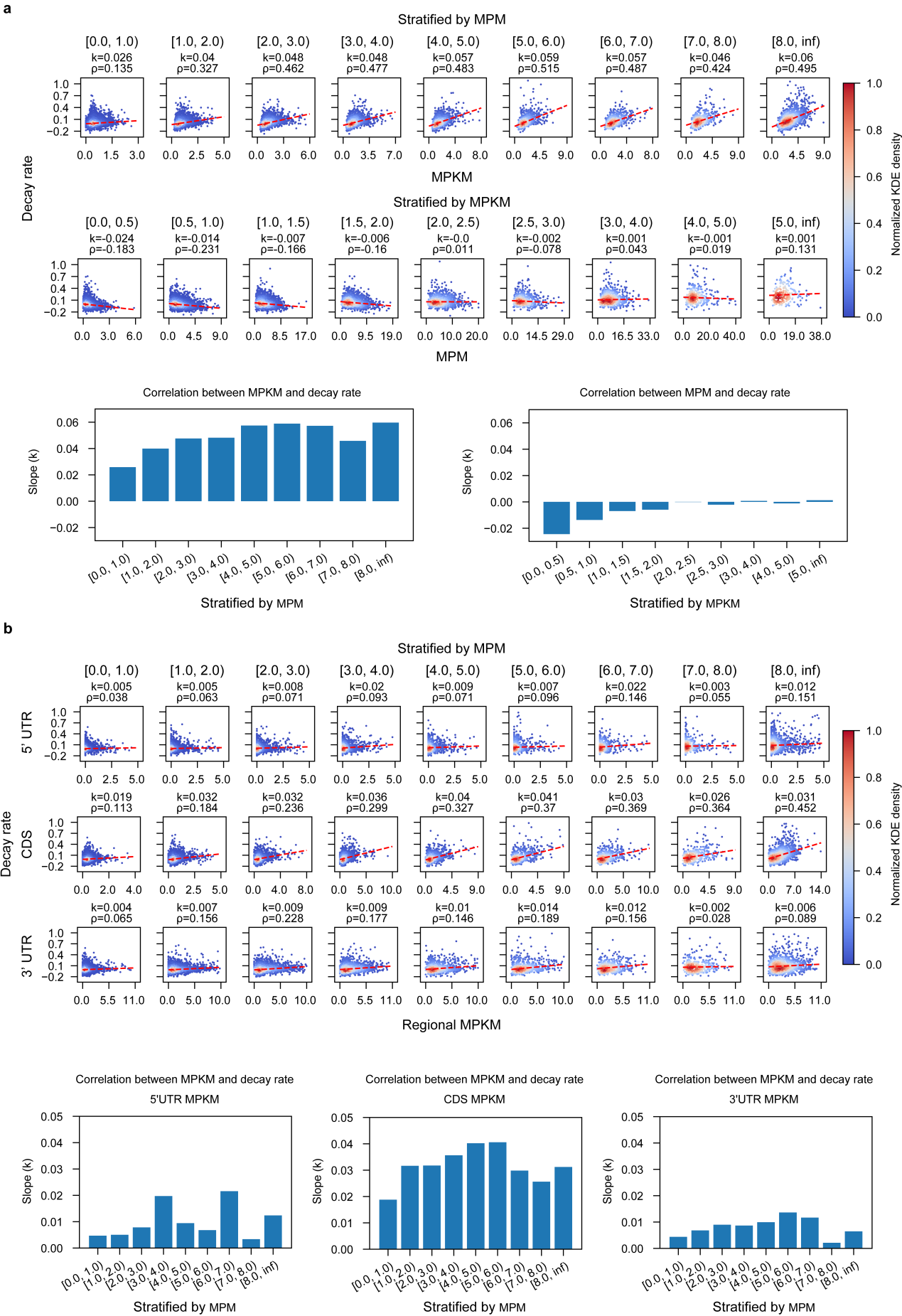
**Supplementary Fig. 7 | Deconvoluting the effects of absolute m6A count and density on RNA decay. a,** Stratified correlation analysis between m6A metrics and RNA decay. Top, scatter plots of decay rate versus m6A density (MPKM), with isoforms stratified by absolute modification count (MPM). Bottom, scatter plots of decay rate versus MPM, with isoforms stratified by their modification density (MPKM). Regression slopes (k) and Spearman’s correlation coefficients (ρ) are indicated for each bin. The bar plots below summarize the regression slopes (k) for each stratified analysis. **b,** Relationship between regional m6A density and RNA decay rate. Scatter plots show the correlation between decay rate and regional MPKM for the 5′ UTR (top), CDS (middle), and 3′ UTR (bottom). All isoforms are stratified based on regional MPM. Regression slopes (k) and Spearman’s correlation coefficients (ρ) are indicated. The bar plots at the bottom summarize the regression slopes (k) for each region across the MPM strata.

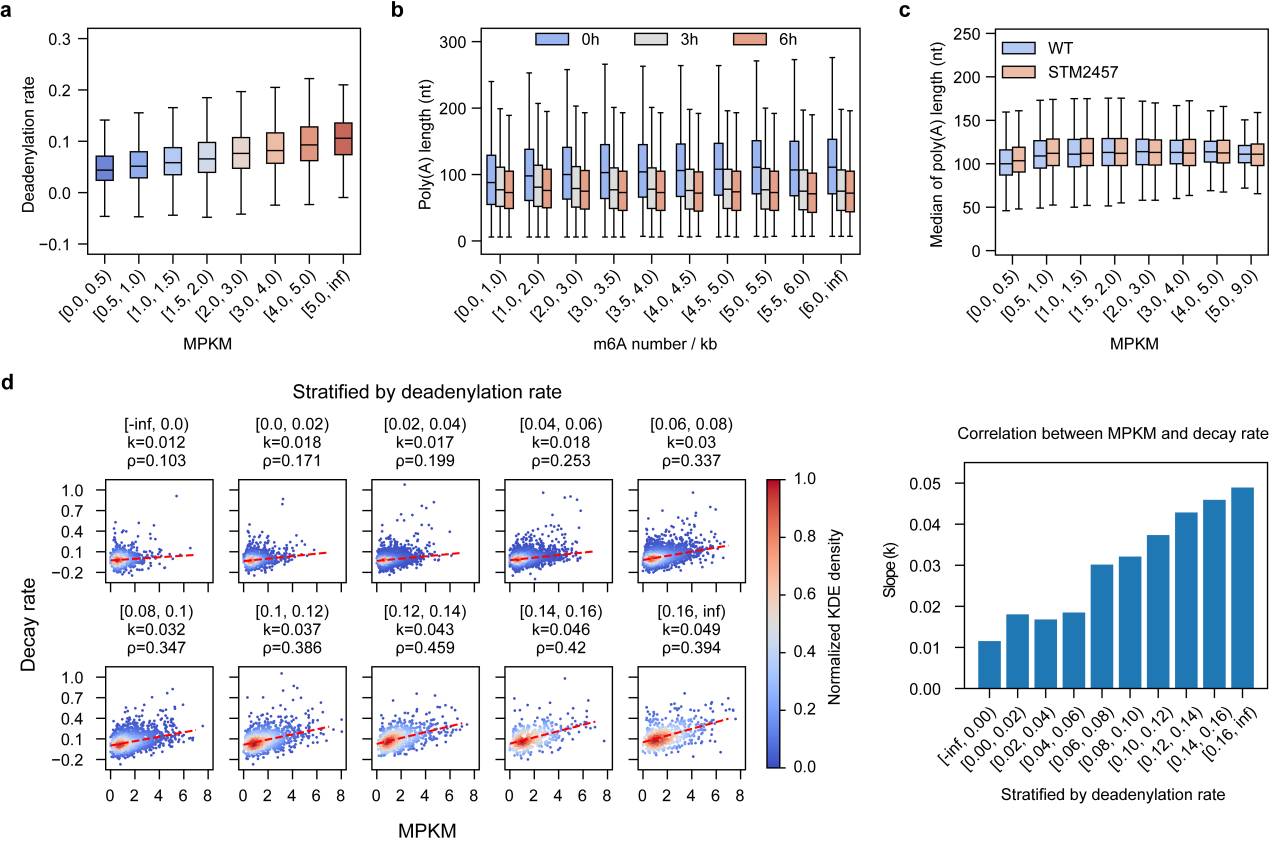

**Supplementary Fig. 8 | Interplay between deadenylation, m6A modification, and gene decay. a,** Deadenylation rates of isoforms stratified by their m6A density (MPKM). **b,** Box plots showing median poly(A) tail length at 0h, 3h, and 6h for isoforms binned by m6A density (m6A number/kb). **c,** Comparison of median poly(A) tail lengths between wild-type (WT) and STM2457-treated cells across isoforms binned by MPKM. **d,** Relationship between RNA decay rate and m6A density (MPKM), stratified by deadenylation rate. Left, scatter plots for isoforms grouped based on their deadenylation rate. Red dashed lines represent robust linear regression fits. Regression slopes (k) and Spearman’s correlation coefficients (ρ) are indicated for each bin. Right, bar plot of regression slopes (k) across deadenylation rate bins. For all box plots, the center line denotes the median, box edges represent the interquartile range (IQR), and whiskers extend to 1.5× the IQR.

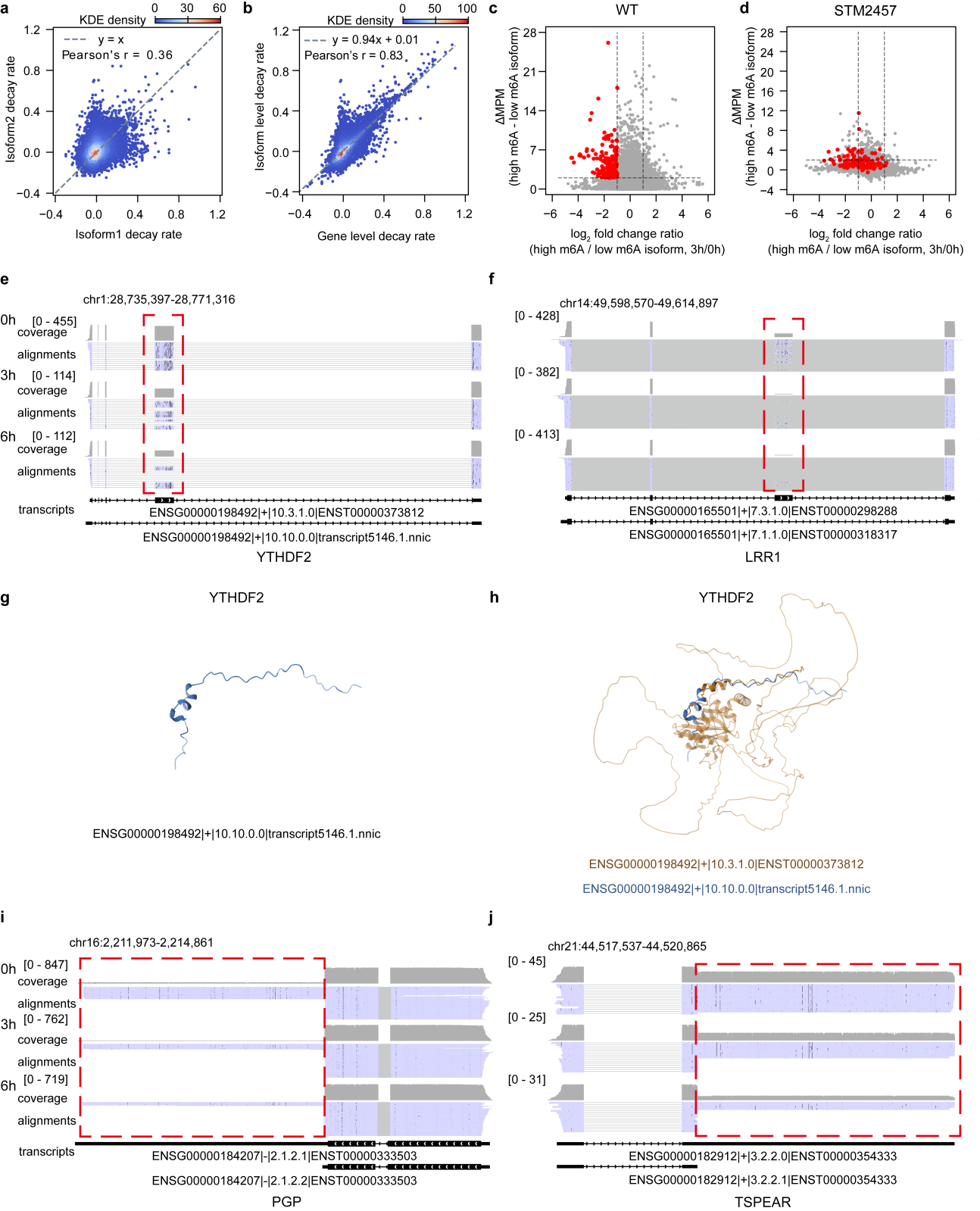
**Supplementary Fig. 9 | RMC-mediated differential degradation and its functional consequences. a,** Comparison of decay rates between pairwise isoforms from the same gene. **b,** Scatter plot comparing isoform-level decay rates to their corresponding gene-level decay rates. **c,** Volcano plot showing the relationship between differential degradation, measured as the log_2_ fold change in relative abundance (at 3h relative to 0h), and the difference in m6A levels (ΔMPM) for high-m6A versus low-m6A isoform pairs in wild-type (WT) cells. Red dots highlight pairs where the isoform with higher m6A abundance exhibits faster degradation (log_2_FC<-1 and ΔMPM>2). **c,** Volcano plot, similar to (**b**), but for cells treated with the METTL3 inhibitor STM2457. **e-f,** Examples of RMC-mediated regulation. IGV snapshots for *YTHDF2* (**e**), and *LRR1* (f) loci in WT cells. Dashed red boxes highlight the isoform-specific regions (RMCs) driving differential stability. **g,** Predicted 3D structure of the truncated *YTHDF2* protein, which results from exon skipping and lacks the m6A-binding YTH domain. **h,** Superimposed 3D structures of the full-length *YTHDF2* (orange) and the truncated product (blue). **i-j,** Examples of RMC-mediated regulation. IGV snapshots for *PGP* (i), *TSPEAR* (j) loci in WT cells. Dashed red boxes highlight the isoform-specific regions (RMCs) driving differential stability.

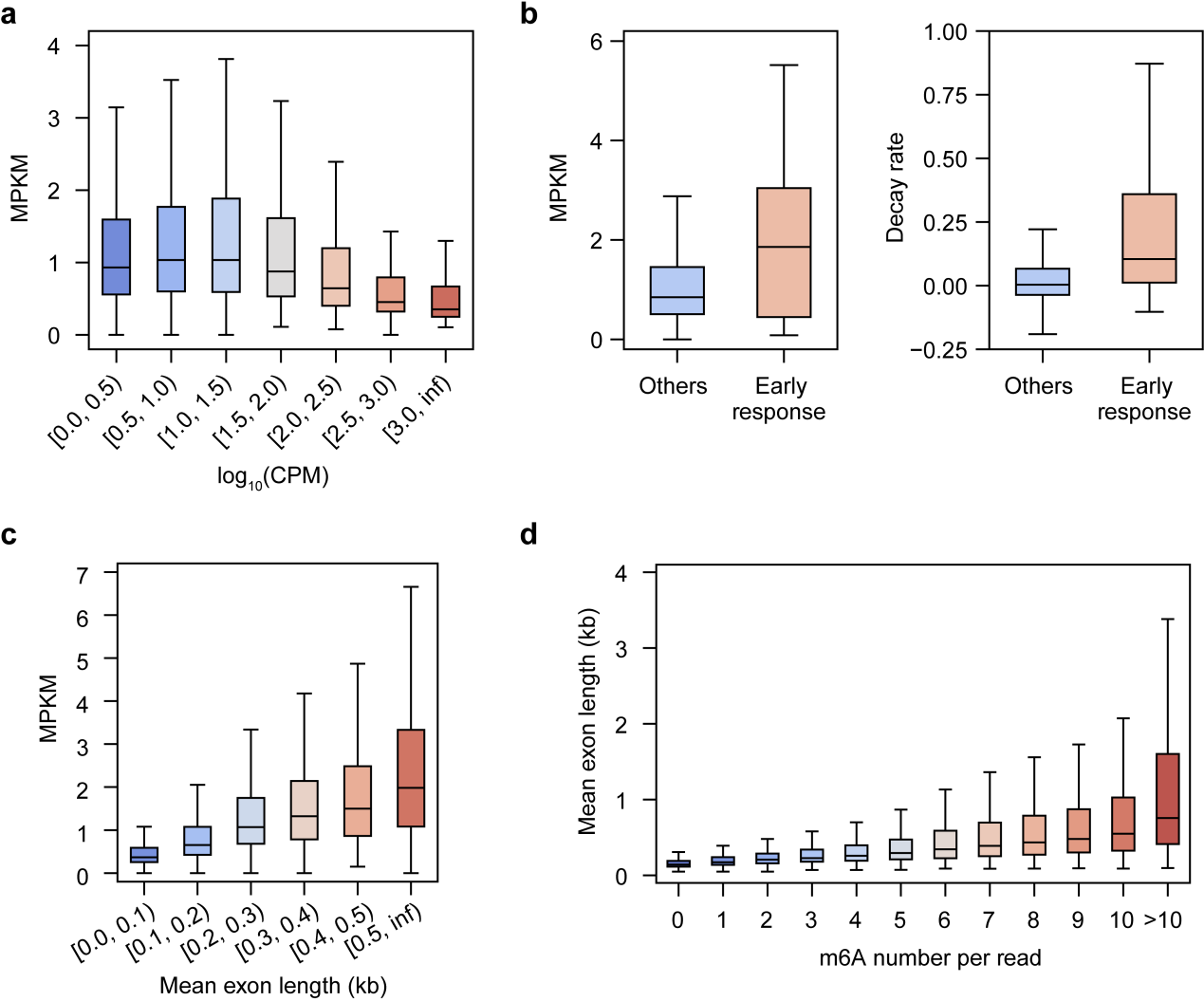

**Supplementary Fig. 10 | Associations between m6A, gene expression, and transcript architecture. a,** Box plot of m6A density (MPKM) for isoforms stratified by their steady-state expression level (log_10_(CPM)). **b**, Comparison of m6A density (MPKM, left) and RNA decay rate (right) between canonical early response genes and all other genes ("Others"). **c,** Box plots of m6A density (MPKM) for isoforms binned by mean exon length (kb). **d,** Box plot showing the mean exon length for reads binned by the number of m6A sites per read. For all box plots, the center line denotes the median, box edges represent the interquartile range (IQR), and whiskers extend to 1.5× the IQR.

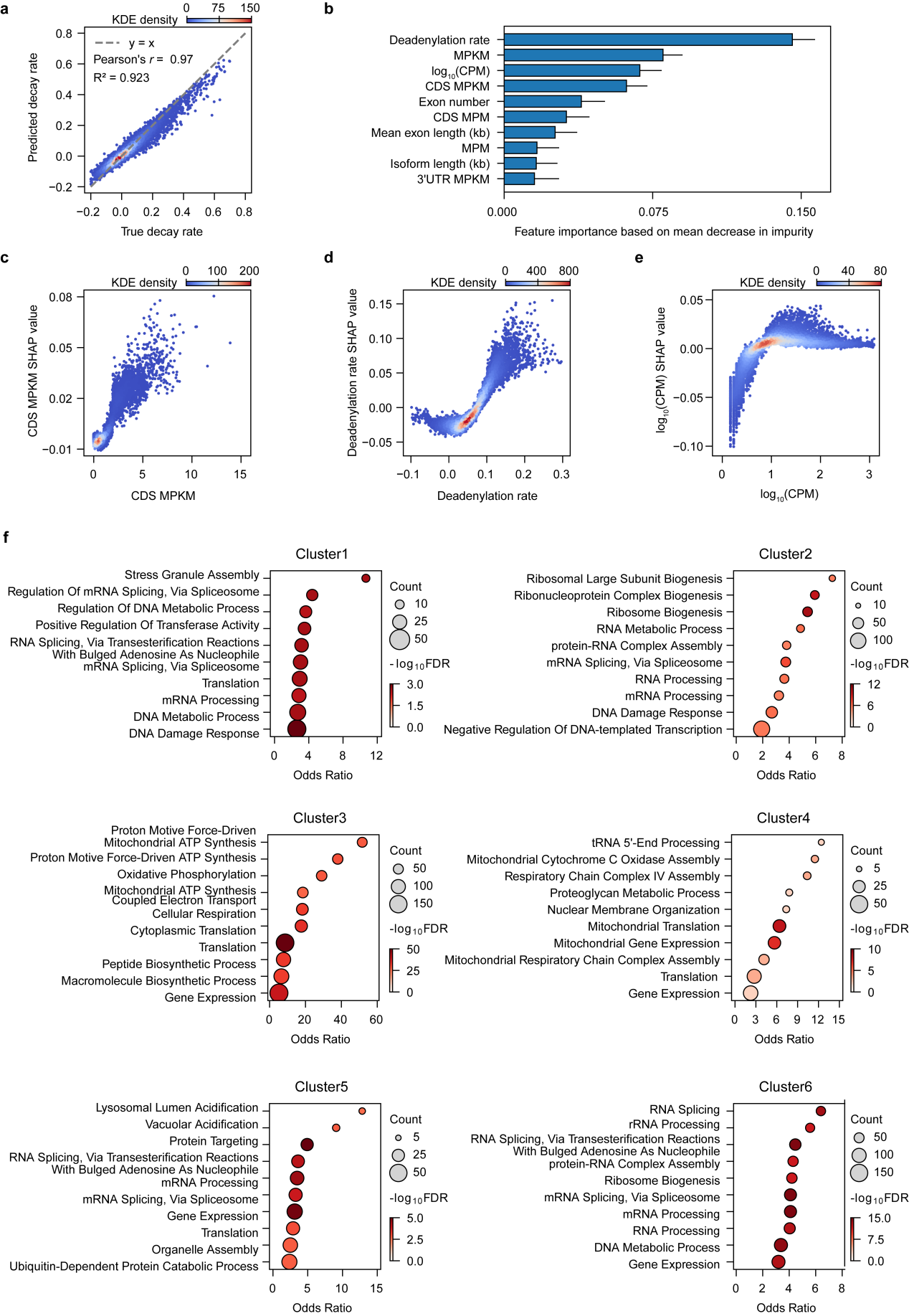

**Supplementary Fig. 11 | Performance evaluation and interpretation of the RNA decay prediction model. a,** Scatter plot comparing decay rates predicted by the Gradient Boosting model versus experimentally determined decay rates on the training set (Pearson's r = 0.97), demonstrating the model's high fitting capability. **b,** Top 10 most important features ranked by impurity-based feature importance derived from the fitted model. **c-e,** SHAP dependence plots for CDS MPKM (**c**), deadenylation rate (**d**), and steady-state expression (log_10_(CPM)) (**e**). Each dot represents an isoform, plotting its feature value against its corresponding SHAP value (impact on model output). **f,** Gene Ontology (GO) enrichment analysis of the six isoform clusters derived from hierarchical clustering of SHAP values (see Fig. 5f). The dot size represents gene count, and the color represents statistical significance (-log_10_FDR).
